# Exapted CRISPR-Cas12f homologs drive RNA-guided transcription

**DOI:** 10.1101/2025.06.10.658865

**Authors:** Florian T. Hoffmann, Tanner Wiegand, Adriana I. Palmieri, Juniper Glass-Klaiber, Renjian Xiao, Stephen Tang, Hoang C. Le, Chance Meers, George D. Lampe, Leifu Chang, Samuel H. Sternberg

**Affiliations:** Department of Biochemistry and Molecular Biophysics, Columbia University, New York, NY, USA; Howard Hughes Medical Institute, Columbia University, New York, NY, USA; Department of Cellular Physiology and Biophysics, Columbia University, New York, NY, USA; Department of Biological Sciences, Purdue University, West Lafayette, IN, USA.; Purdue Institute for Cancer Research, Purdue University, West Lafayette, IN, USA; Department of Medicine, Stanford University, Stanford, CA, USA; Department of Biochemistry, Vanderbilt University, Nashville, TN, USA

## Abstract

Bacterial transcription initiation is a tightly regulated process that canonically relies on sequence-specific promoter recognition by dedicated sigma (σ) factors, leading to functional DNA engagement by RNA polymerase (RNAP)^1^. Although the seven σ factors in *E. coli* have been extensively characterized^2^, Bacteroidetes species encode dozens of specialized, extracytoplasmic function σ factors (σ^E^) whose precise roles are unknown, pointing to additional layers of regulatory potential^3^. Here we uncover an unprecedented mechanism of RNA-guided gene activation involving the coordinated action of σ^E^ factor in complex with nuclease-dead Cas12f (dCas12f). We screened a large set of genetically-linked dCas12f and σ^E^ homologs in *E. coli* using RIP-seq and ChIP-seq experiments, revealing systems that exhibited robust guide RNA enrichment and DNA target binding with a minimal 5ʹ-G target-adjacent motif (TAM). Recruitment of σ^E^ was dependent on dCas12f and guide RNA (gRNA), suggesting direct protein-protein interactions, and co-expression experiments demonstrated that the dCas12f-gRNA-σ^E^ ternary complex was competent for programmable recruitment of the RNAP holoenzyme. Remarkably, dCas12f-RNA-σ^E^ complexes drove potent gene expression in the absence of any requisite promoter motifs, with *de novo* transcription start sites defined exclusively by the relative distance from the dCas12f-mediated R-loop. Our findings highlight a new paradigm of RNA-guided transcription (RGT) that embodies natural features reminiscent of CRISPRa technology developed by humans^4,5^.

## INTRODUCTION

Canonical CRISPR-Cas systems provide adaptive immunity in bacteria and archaea by eliminating foreign nucleic acids from invading bacteriophages and plasmids^6^. Within the broad CRISPR-Cas diversity, Cas9 and Cas12 represent archetypal single-effector proteins that target and cleave double-stranded DNA (dsDNA) substrates recognized via base-pairing to a guide RNA (gRNA) and the protospacer-adjacent motif (PAM)^7,8^. Both proteins are ancestrally related to homologous, transposon-encoded nucleases within the TnpB superfamily, which similarly use gRNAs to target dsDNA for cleavage based on RNADNA complementary and target-adjacent motif (TAM) recognition, thereby promoting selfish transposon spread^9–12^. The recent discovery of RNA-guided transposases encoded by IS110-family transposons^13,14^, as well as their striking resemblance to eukaryotic homologs comprising Nop5 domains^15^, establishes the pervasive degree to which mobile, noncoding RNAs were exapted for essential functions ranging from RNA methylation and RNA splicing to RNA-guided DNA recognition.

TnpB enzymes, whose eukaryotic homologs are known as Fanzors^10,16^, are one of the most abundant prokaryotic proteins^17^. Although they primarily expanded in copy number via transposon-mediated horizontal gene transfer, many TnpB subfamilies exhibit low mobility scores and unusual genomic neighborhoods indicative of novel function^17^; in some cases, their RuvC nuclease domains harbor active site mutations expected to abolish DNA cleavage behavior^17,18^. We recently reported numerous instances of TnpB-like nuclease dead repressors (TldR), whose RNA-guided DNA binding activities direct potent gene silencing that facilitates flagellar remodeling of the host (**Fig. 1a**)^18^. This finding reframes CRISPR-Cas as just one pathway among countless TnpB domestication trajectories explored across evolutionary timescales, while also highlighting the recurrent co-option of RNA-guided DNA binding proteins for transcriptional reprogramming. Even within CRISPR-Cas systems, Cas9 nucleases (Type II) are repurposed as RNA-guided gene repressors for autoregulatory and virulence control^19–21^; Cas12 proteins (Type V) sometimes mediate adaptive immunity by blocking transcription rather than cleaving DNA^22,23^; and Cascade complexes (Type I) can silence toxin RNAs to promote their selfish retention^24^. Collectively, these naturally occurring examples resemble the CRISPR interference (CRISPRi) technologies that have been developed for RNA-guided transcriptional silencing^25–27^.

**Figure 1.**
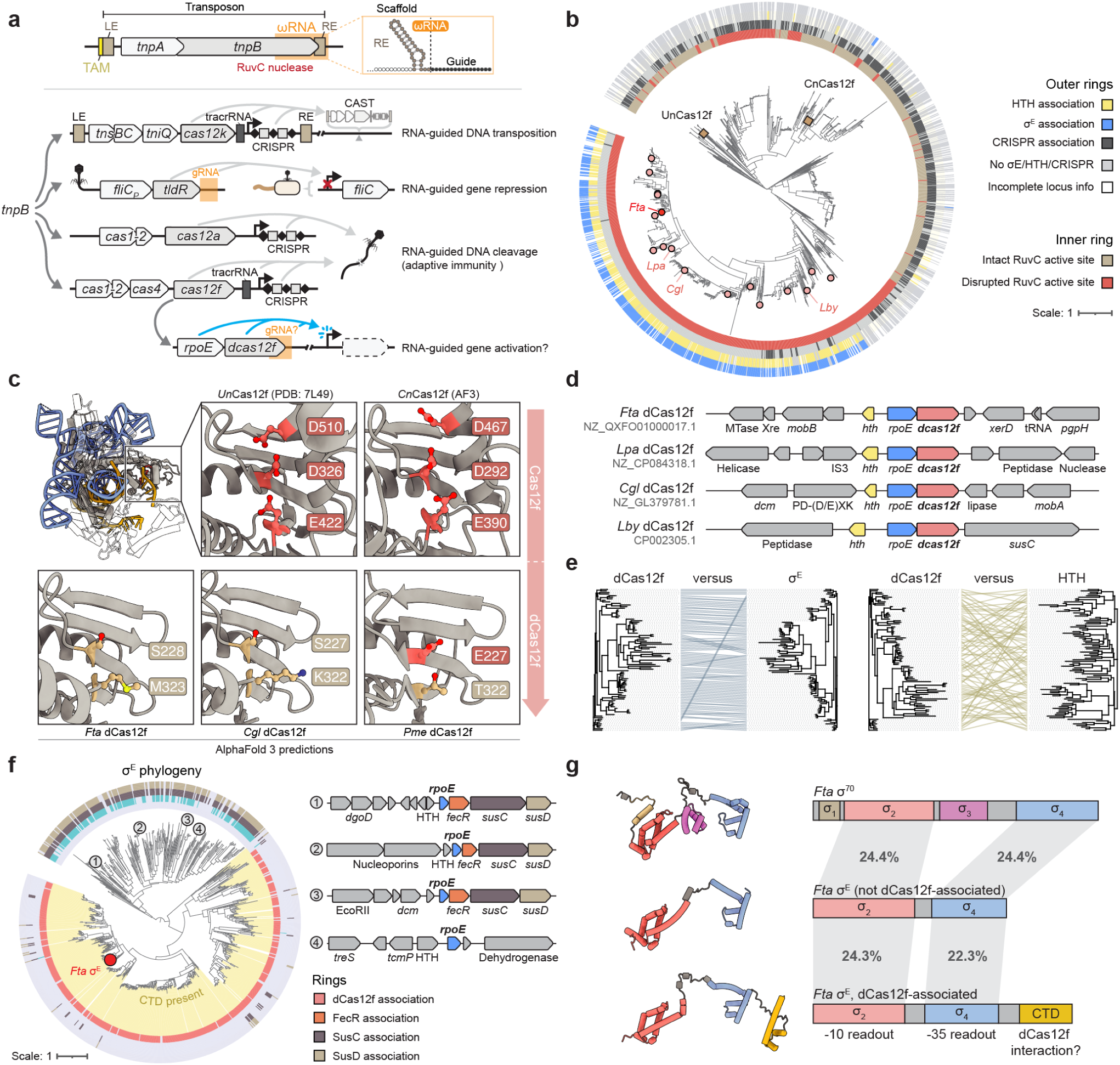
Nuclease-dead Cas12f homologs are genetically associated with atypical σ factor genes. **a,** Schematic depicting exaptation events of transposon-encoded *tnpB* nuclease genes for novel, RNA-guided functions. Primordial *tnpB* homologs (top) are encoded within the left end (LE) and right end (RE) boundaries of bacterial transposons alongside the *tnpA* transposase, where they function together with ωRNA guides to promote transposon retention. Recurrent *tnpB* domestication gave rise to diverse CRISPR-associated effectors such as Cas12a–Cas12f, CRISPR-associated transposon (CAST) proteins such as Cas12k, and RNA-guided transcriptional repressors such as TldR (middle). Nuclease-dead dCas12f proteins (bottom) are proposed to function with gRNAs and σ^E^ factors (encoded by *rpoE*) to drive gene activation. **b,** Phylogenetic tree of Cas12f homologs, showing progressive RuvC nuclease domain inactivation (inner ring, grey/red), loss of CRISPR array association, and gain in HTH and σ^E^ (*rpoE* gene) association. Biochemically characterized Cas12f nucleases (red squares) and candidate nuclease-dead dCas12f homologs (red circles) are indicated; the homolog from *Flagellimonas taeanensis* (*Fta*) was selected for detailed characterization. **c,** Magnified view of the RuvC nuclease domain from AlphaFold 3 structure predictions of select dCas12f homologs from **b** (bottom), compared to structurally characterized Cas12f nucleases (top). Active site residues and their corresponding naturally occurring mutations are labeled. **d,** Representative genomic neighborhoods of select dCas12f homologs from **b**, with labeled genomic accession IDs. **e,** Comparison of phylogenetic trees for select dCas12f, σ^E^, and HTH homologs. This analysis reveals that *dCas12f* and *rpoE* genes exhibit a strong genetic linkage, whereas *dCas12f* and *hth* genes do not. **f**, Phylogenetic tree of σ^E^ homologs, showing progressive loss of FecR and SusCD associations (outer rings) and gain of dCas12f association (inner ring, red); the highlighted yellow sector represents the concomitant gain of the C-terminal domain (CTD). Representative genomic neighborhoods for the indicated leaves (1–4) are shown at right. **g,** AlphaFold 3 structure predictions (left) and domain annotations (right) for three σ factors from *F. taeanensis*: σ^70^ (*rpoD* gene), a representative σ^E^ homolog (*rpoE*) that is non-dCas12f-associated, and the dCas12f-associated σ^E^ (*rpoE*) from **d**. Each protein shares homologous σ and σ domains, but the dCas12f-associated σ^E^ uniquely exhibits an additional CTD that may interact with dCas12f.

Have TnpB or CRISPR-associated proteins also been coopted for transcriptional *activation*, akin to CRISPRa technologies developed by humans^4,5,28,29^? We became intrigued by unusual Cas12f genes, described in a recent report^17^, which harbor inactivating RuvC mutations and are encoded adjacent to sigma (σ) factor genes, suggesting a potential role in transcription initiation. In bacteria, σ factor is an essential subunit of the RNA polymerase (RNAP) holoenzyme that imparts sequence-specific promoter DNA recognition^1^. Promoter interactions without σ are insufficient to initiate transcription, underpinning the central role of σ in gene expression^30,31^. σ factor genes have been comprehensively studied in *E. coli*, where seven homologs control transcriptional programs associated with normal growth, stress adaptation, nutrient availability, and motility control^32,33^, but their numbers are massively expanded in other bacteria. Group 4 σ factors in particular, also known as the extracytoplasmic function (ECF) family because of their roles in envelope stress response and metabolite uptake^34–38^, are highly abundant in Bacteroidetes^3^ and encompass the specific subtype found adjacent to Cas12f genes. We therefore hypothesized that these unusual operons, encoding nuclease-dead Cas12f (hereafter dCas12f) and ECF-like sigma (σ^E^) factors, might encode novel machinery to perform RNA-guided transcription of target genes.

Here, we report the discovery and characterization of a natural bacterial pathway for programmable gene activation. After bioinformatically exploring dCas12f-σ^E^ systems, we reconstituted their activity in *E. coli*, demonstrating that dCas12f-gRNA complexes recognize target sites through RNA-DNA base-pairing and a minimal TAM. Target binding triggers recruitment of σ^E^-RNAP, driving transcription initiation at a fixed distance downstream of the target site. In native strains, dCas12f-σ^E^ systems control the expression of conserved transmembrane transport systems implicated in nutrient import and utilization, and are likely modulated by associated transcriptional regulators. Collectively, our findings uncover another unexpected evolutionary link between CRISPR-Cas systems and transcriptional control, expanding the functional repertoire of RNA-guided mechanisms in bacteria.

## RESULTS

### Genetic association between Cas12f and σ^E^

We developed a bioinformatics pipeline to identify Cas12-like proteins hypothesized to lack DNA cleavage activity based on the presence of characteristic RuvC nuclease domain mutations, building from our previous TnpB work^18^. This analysis uncovered a large group of inactivated effectors closely related to Cas12f, of which the majority lacked a CRISPR association but exhibited a strong association with σ^E^ factors^34,39^ and helix-turn-helix (HTH) domain proteins (**Fig. 1b**), paralleling recent observations by Zhang and colleagues^17^. We also identified a distinct cluster of putative nuclease-dead Cas12f (dCas12f) proteins encoded within the vicinity of CRISPR arrays, whose predicted gRNAs sometimes target the promoter region of efflux system expression cassettes, suggesting a potential role in transcriptional repression (**Extended Data Fig. 1a,b**). Compared to nuclease-active Cas12f^40^, dCas12f proteins harbor C-terminal truncations that eliminate part of the RuvC active site, but AlphaFold 3 structural models suggested they nonetheless maintain a Cas12f-like fold (**Fig. 1c**).

The strong co-localization and operonic arrangement of genes encoding dCas12f (*dcas12f*) and σ^E^ (*rpoE*; **Fig. 1d**), suggested an intriguing potential coordination between RNA-guided DNA binding and transcription initiation. The association with HTH proteins (encoded by *hth*), whose functions also generally involve transcriptional control^41^, was more enigmatic, and phylogenetic analyses indicated a greater degree of HTH diversification and/or recombination within *rpoE-dcas12f* loci, compared to the clear pattern of dCas12f and σ^E^ co-evolution (**Fig. 1e, Extended Data Fig. 1c**).

Further analysis of σ^E^ phylogeny provided additional clues supporting its predicted synergy with dCas12f (**Fig. 1f**). Canonical σ^E^ factors are encoded next to anti-σ factors such as FecR, which reside in the inner membrane and sequester σ^E^ until sensing of an extracellular stimulus activates FecR cleavage and concomitant σ^E^ release, leading to target gene transcription^42^. We observed genetic architectures consistent with this paradigm for σ^E^ clades that were distantly related to dCas12f-σ^E^, with *rpoE-fecR* loci furthermore associated with *susCD* genes involved in transmembrane transport (**Fig. 1f**). In contrast, *rpoE-dcas12f* loci consistently lacked FecR and SusCD associations, suggesting distinct biological functions. When we analyzed the σ^E^ domain composition more closely, we noticed that dCas12f-associated homologs possessed a unique C-terminal domain (CTD) extension, as was reported previously^17^, which we hypothesized could mediate dCas12f interactions. This domain was absent from nearly all non-dCas12f-associated homologs (**Fig. 1f,g**)^17^. The only notable exception was a comparatively rare clade comprising σ^E^ homologs associated with unusual restriction endonuclease–like proteins also predicted to be nuclease inactive (**Extended Data Fig. 1d**).

Collectively, these bioinformatics observations strongly suggested a novel role for dCas12f proteins involving transcriptional regulation by σ^E^ and/or HTH, unrelated to CRISPR-Cas immunity. We were especially interested in testing the hypothesis that RNA-guided DNA targeting by dCas12f might lead to functional recruitment of σ^E^ and RNA polymerase.

### dCas12f exhibits RNA-guided DNA binding

To experimentally investigate dCas12f function, we devised a heterologous expression strategy in *E. coli* that would reveal gRNA and target DNA binding motifs via RNA and chromatin immunoprecipitation sequencing (RIP-seq, ChIP-seq), respectively (**Fig. 2a**)^18,43^. A previous study described ωRNA-like arrays associated with dCas12f-σ^E^ systems, though without clearly defining the identity of mature gRNAs^17^. We selected 16 phylogenetically diverse dCas12f-σ^E^ systems (**Fig. 1b, Extended Data Fig. 2**) and cloned both genes on a single effector plasmid, alongside *hth* and candidate regions that we hypothesized would contain gRNAs (**Fig. 2a**). The resulting RIP-seq data revealed highly enriched gRNAs encoded immediately downstream of *dcas12f*, with clearly demarcated ends and average lengths of 85–105 nt (**Fig. 2b,c, Extended Data Fig. 3a,b**). dCas12f homologs from *Flagellimonas taeanensis* (*Fta*) and *Leadbetterella byssophila* (*Lby*) associated with a single gRNA, whereas *Leeuwenhoekiella palythoae* (*Lpa*) and *Paenimyroides ummariense* (*Pum*) homologs enriched multiple tandemly encoded gRNAs with similar sequence (**Fig. 2c, Extended Data Fig. 3b,c**).

**Figure 2.**
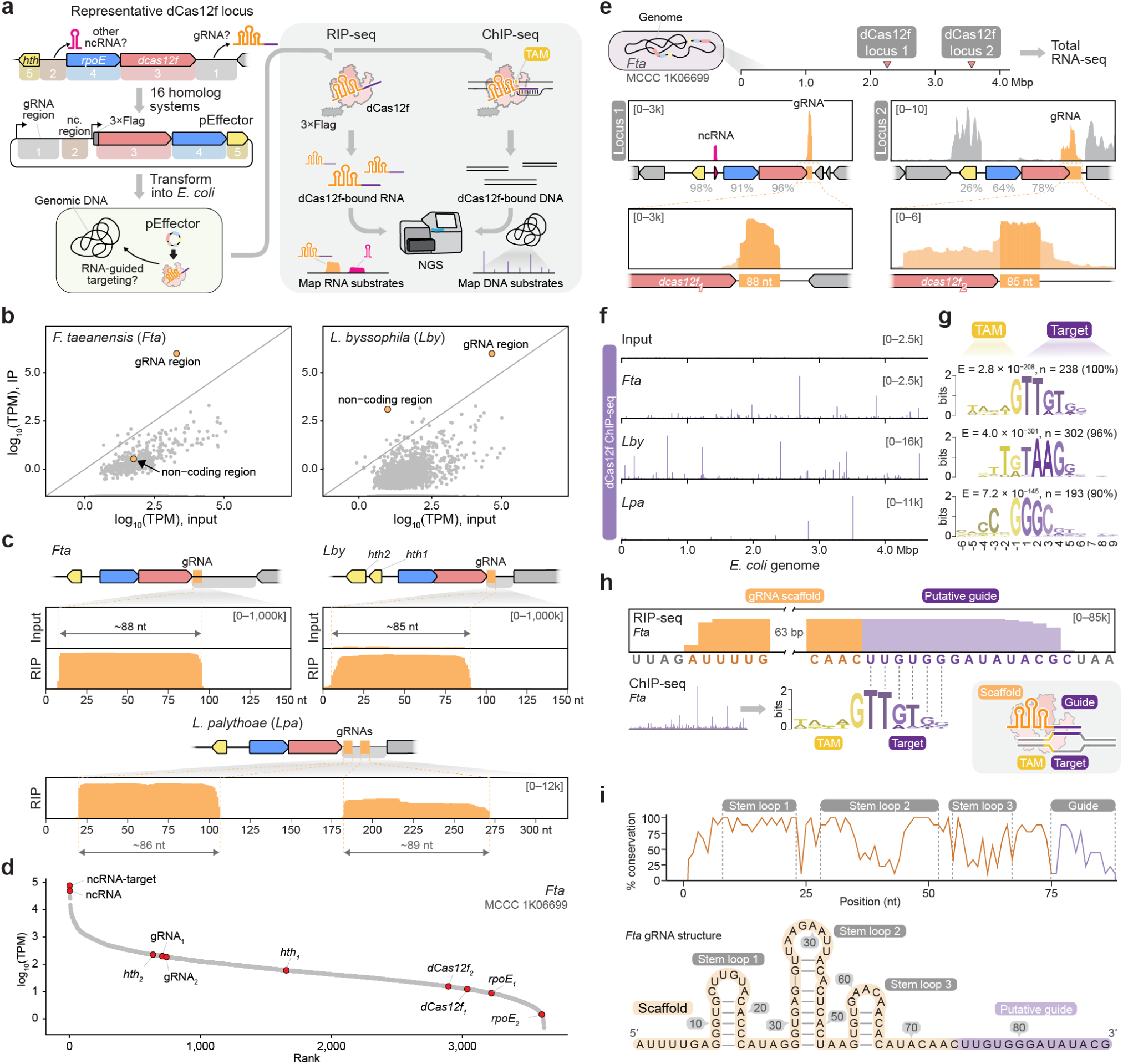
Experimental discovery of guide RNA and target DNA substrates of dCas12f. **a,** Vector design (left) and experimental workflow (right) to identify bound gRNAs and target DNA sites by RIP-seq and ChIP-seq, respectively. **b,** Representative RIP-seq data for *Fta* (left) and *Lby* (right) dCas12f, with the scatter plot comparing log-transformed transcripts per million (TPM) in the IP fraction (y-axis) versus the input fraction (x-axis). **c,** RIP-seq coverage plots comparing the input and IP fraction for three dCas12f homologs, with the corresponding genomic neighborhoods schematized above. dCas12f associates with a well-defined gRNA that is 85–89 nucleotides (nt) in length, encoded immediately downstream of the *dcas12f* gene. The *Lpa* homolog co-purifies with two gRNAs encoded in tandem (bottom). **d,** Ranked transcript abundance, plotted as log-transformed TPM, from RNA-seq data of an *F. taeanensis* strain that encodes two dCas12f-σ^E^ systems. Key genes and RNAs are labeled. **e,** RNA-seq coverage plots from data shown in **d,** mapped onto the two *rpoE–dcas12f* loci; magnified insets are shown below. The percent sequence identities between these proteins and corresponding *Fta* homologs tested heterologously in *E. coli* are shown. **f,** Genome-wide representation of ChIP-seq data for the indicated dCas12f homologs analyzed in **c** (purple), compared to the input control (top). Coverage is shown as counts per million (CPM), normalized to the highest peak in the targeting sample. **g,** Binding events were analyzed by MEME-ChIP, revealing consensus motifs that correspond to the TAM and gRNA-matching target DNA sequence within the seed. E, E-value significance; n, number of peaks contributing to the motif. **h,** Schematic demonstrating how a comparison of RIP-seq and ChIP-seq motif data results in high-confidence identification of the TAM and target DNA-gRNA base-pairing, consistent with the expected R-loop complex (bottom right). **i,** Sequence (top) and secondary structure (bottom) of the *Fta* dCas12f-associated gRNA from RIP-seq data in **b,c**, with labeled stem loop and guide features. The graph shows sequence conservation across an alignment of nine gRNAs identified in RIP- or RNA-seq experiments.

To further corroborate these results, we obtained multiple *Fta* strains, developed culturing conditions (**Methods**), and performed genome and RNA sequencing (**Extended Data Fig. 4a,b**). *Fta* strain 4759 encodes two *rpoE*-*dcas12f* loci (**Extended Data Fig. 4b,c**), and RNA-seq analysis demonstrated that gRNAs from both were highly expressed and closely matched RIP-seq results (**Fig. 2d,e**). The *rpoE* and *dcas12f* genes were themselves lowly expressed (**Fig. 2d**), suggesting that induction relies on a yet unidentified physiological state. Intriguingly, we also observed an abundant non-coding RNA (ncRNA) within the intergenic region between *hth* and *rpoE*, as well as a similar ncRNA adjacent to an *hth* homolog elsewhere in the genome (**Fig. 2d,e, Extended Data Fig. 4d**) suggesting potential regulatory roles (see below).

Next, we performed ChIP-seq with the same 16 systems, hypothesizing that off-target sites resembling the gRNA would be promiscuously bound by dCas12f, enabling unambiguous identification of the gRNA guide sequence and target adjacent motif (TAM). The resulting data revealed hundreds of highly enriched regions for dCas12f homologs whose gRNAs were readily identified by RIP-seq (**Fig. 2f, Extended Data Fig. 5a**), and subsequent analyses uncovered conserved motifs common to the majority of peaks within each dataset (**Fig. 2g, Extended Data Fig. 5b**). By integrating these motif results with RIP-seq data and RNA structural analyses, we confidently deduced the TAM and a 14–16-nt guide sequence for 7 distinct dCas12f systems (**Fig. 2h, Extended Data Fig. 3a-c, 5b**). Complementarity was most critical within a short TAM-proximal region recognized by the gRNA seed sequence^12,40^, and notably, the predicted TAM for the *Fta* dCas12f homolog was just a single 5’-G nucleotide (**Fig. 2h**).

Our analyses failed to identify tracrRNA-like species, in contrast to Cas12f1 homologs from Type V-F1 CRISPR-Cas systems^6,44,45^. Instead, our empirically identified gRNAs revealed a stem-loop architecture more reminiscent of TnpB-associated ωRNAs, with a guide sequence positioned at the 3’ end (**Fig. 2i**)^12,17,18,46^. We therefore concluded that dCas12f homologs function with single guide RNAs comprising both a scaffold and guide portion, and hypothesized that they might have evolved from an ancestral state more akin to CRISPR arrays. In agreement with this notion, we identified an *rpoE-dcas12f* locus from *Sphingobacterium* that appears to encode full-length gRNAs closely resembling the repeat sequences found in CRISPR arrays interspersed throughout the same locus (**Extended Data Fig. 3d**). This observation suggests the intriguing possibility that dCas12f-associated gRNAs emerged from the repeat-spacer sequence of CRISPR arrays themselves.

### Identification of dCas12f-gRNA targets

We next sought to identify putative genomic targets of dCas12f, based on the hypothesis that RNA-guided σ^E^ recruitment would lead to RNAP binding and transcription initiation. Strikingly, when we searched the *Fta* genome for gRNA matches, we immediately identified candidates within the large intergenic region upstream of a *susCD* operon implicated in nutrient import and utilization (**Fig. 3a,b**). Both sites were flanked by a 5’-G TAM, exhibited 13-14 matching base pairs with the predicted guide, and were positioned to potentially serve promoter-like functions (**Fig. 3b**). Intriguingly, the very same genomic region encoded one of the *hth*-associated ncRNAs noted earlier, which overlapped precisely with the gRNA 5’ edge and exhibited the same strandedness (**Fig. 3a,b**).

**Figure 3.**
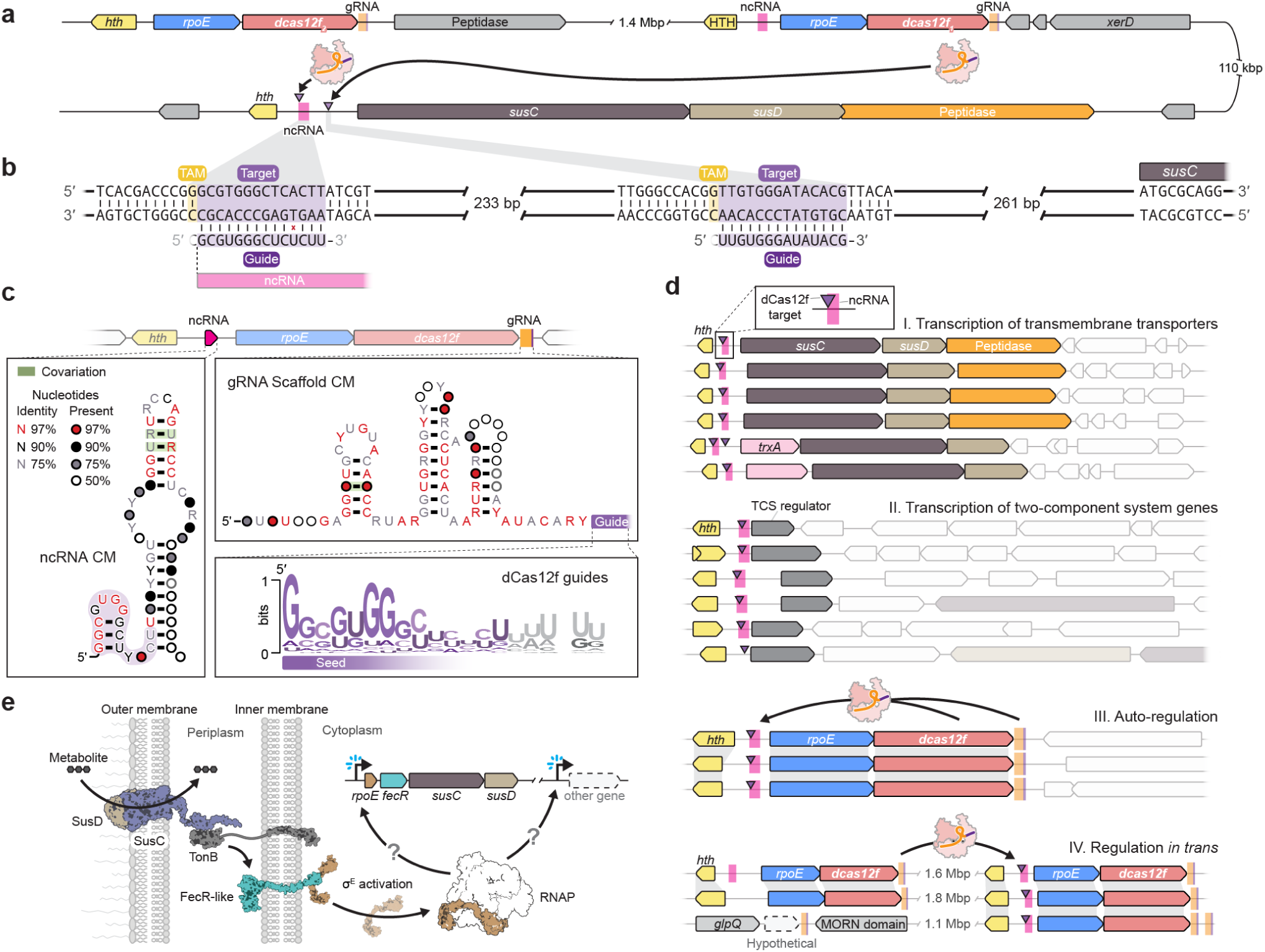
dCas12f-associated gRNAs target conserved ncRNA loci and regulate diverse gene expression programs. **a,** Schematic of genomic sites (1, 2) targeted by gRNAs associated with two *dcas12f* operons in the *Fta* strain from Fig. 2d. The gRNA.2-matching target site (purple triangle) precisely overlaps the site of an HTH-associated ncRNA (magenta rectangle), which itself closely resembles a similar ncRNA associated with the *hth* gene in the dCas12f.1 operon. **b,** Schematic of RNADNA complementarity for targets 1 and 2, with the TAM and guide sequences labeled; the guide sequence at target 1 overlaps the 5′ end of the HTH-associated ncRNA. **c,** Covariance models (CM) of the *hth*-associated ncRNA (left) and gRNA scaffold (right), with conserved and covarying positions represented as shown in the legend. The WebLogo depicts conservation of the guide sequence, whose 5′ seed sequence closely resembles the 5′ end of the *hth*-associated ncRNA (shaded purple region). **d**, Schematic showing four categories of genetic loci targeted by dCas12f-associated gRNAs, with predicted target sites (purple triangles) and predicted *hth*-associated ncRNAs (magenta rectangles) indicated. The most frequent targets are transmembrane transporters (I) and two-component system (TCS) regulator genes (II), but gRNAs also likely control expression of the *rpoEdcas12f* operon itself via auto-regulation *in cis* or regulation *in trans* (III and IV). **e,** Model of canonical σ^E^-mediated transcriptional activation of *susCD* and/or other genes to regulate outer membrane transport systems. SusC and SusD mediate the uptake of metabolites across the outer membrane in coordination with TonB, whose activities are signaled through a FecR-like protein to activate σ^E^ and drive transcription at specific promoters.

To comprehensively explore dCas12f targeting, we systematized the search by building gRNA and ncRNA covariance models (**Fig. 3c**) and globally interrogating genomes with *rpoE-dcas12f* loci to identify their putative genomic targets (**Extended Data Fig. 6a,b**). This analysis revealed four categories of predicted dCas12f function, which were consistent with potential roles in transcription initiation (**Extended Data Fig. 6c**). Most of the queried genomes harbored RNA-guided DNA targets upstream of *susCD* operons involved in transmembrane trafficking (category I) (**Fig. 3d, Extended Data Fig. 6d**), similar to the *Fta* genome (**Fig. 3a,b**). We identified other targets upstream of loci encoding two component systems, with roles in cell signaling (category II) (**Fig. 3d, Extended Data Fig. 6d**). Surprisingly, we also observed gRNA-matching target sites upstream of *rpoE-dcas12f* loci themselves, with gRNAs encoded either immediately downstream of the locus, suggesting positive auto-regulatory feedback *in cis* (category III), or elsewhere in the genome, suggesting a more complex *trans*-acting regulatory network (category IV) (**Fig. 3d**). In some cases, we identified gRNAs that putatively target up to seven distinct loci spanning multiple functional categories (**Extended Data Fig. 6e**), suggesting that *rpoE-dcas12f* loci are responsible for the coordinated regulation of entire gene programs spread across the genome.

The conspicuous presence of gRNA-matching target sites upstream of protein-coding genes strongly corroborated our hypothesis that R-loop formation by dCas12f-gRNA complexes could result in σ^E^-RNAP recruitment, thereby leading to transcription initiation. Furthermore, the frequent presence of targets upstream of TonB-linked SusCD transporters strongly implicates these systems in regulating nutrient uptake, and TonB-linked transporters are indeed canonically regulated by σ^E^ factors, which increase expression when the extracellular nutrient concentration escalates (**Fig. 3e**)^47,48^. Our observation that some *rpoE-dcas12f* loci are natively encoded adjacent to *susCD* operons (**Fig. 1d**), and that non-dCas12f-associated *rpoE* genes are typically encoded near similar *susCD* gene clusters (**Fig. 1f**), strongly supports this functional link. We even detected instances in which CRISPR arrays encoded adjacent to *rpoE-dcas12f* loci harbor conventional spacers that target *susCD* operons, consistent with the hypothesis that *susCD*-specific spacers may have been integrated prior to the emergence of chimeric single gRNAs (**Extended Data Fig. 6f**).

Intriguingly, we consistently predicted *hth*-associated ncRNAs that precisely overlap with gRNA-matching target sites across each of these major categories (**Fig. 3c,d, Extended Data Fig. 6b,e**). Moreover, these gRNAs and ncRNAs are encoded on the same strand, ruling out a role for sense-antisense transcriptional interference. Although the biological roles of the ncRNA remain unknown, as do the promoter elements controlling their expression, we hypothesize that *hth*-associated ncRNAs function as negative regulators of the same genes acted upon by dCas12f-σ^E^ systems, similar to the role of anti-σ factors regulating canonical σ^E^.

### dCas12f-gRNA recruits σ^E^ to target DNA

Having identified the native guides and TAMs for dCas12f-σ^E^, we investigated targeting by reprogramming the gRNA and performing ChIP-seq. Focusing on the *Fta* system, we designed five guides to target sites tiled across the *E. coli* genome, with no selection criteria other than a 5’-G TAM (**Fig. 4a**). Strong enrichment at all five targets confirmed that the minimal TAM and RNA–DNA complementarity were sufficient for dCas12f binding (**Fig. 4a**), and additional dCas12f homologs exhibited similar activities (**Extended Data Fig. 7a**). To determine whether dCas12f recruits σ^E^, we relocated the 3×Flag tag to σ^E^ and performed similar ChIP-seq experiments, which revealed σ^E^ occupancy at the same target sites (**Fig. 4b**). Control experiments further demonstrated that dCas12f and gRNA were necessary and sufficient for target-specific binding of dCas12f, but that σ^E^, HTH, and the *hth*-associated ncRNA were dispensable (**Fig. 4c**).

**Figure 4.**
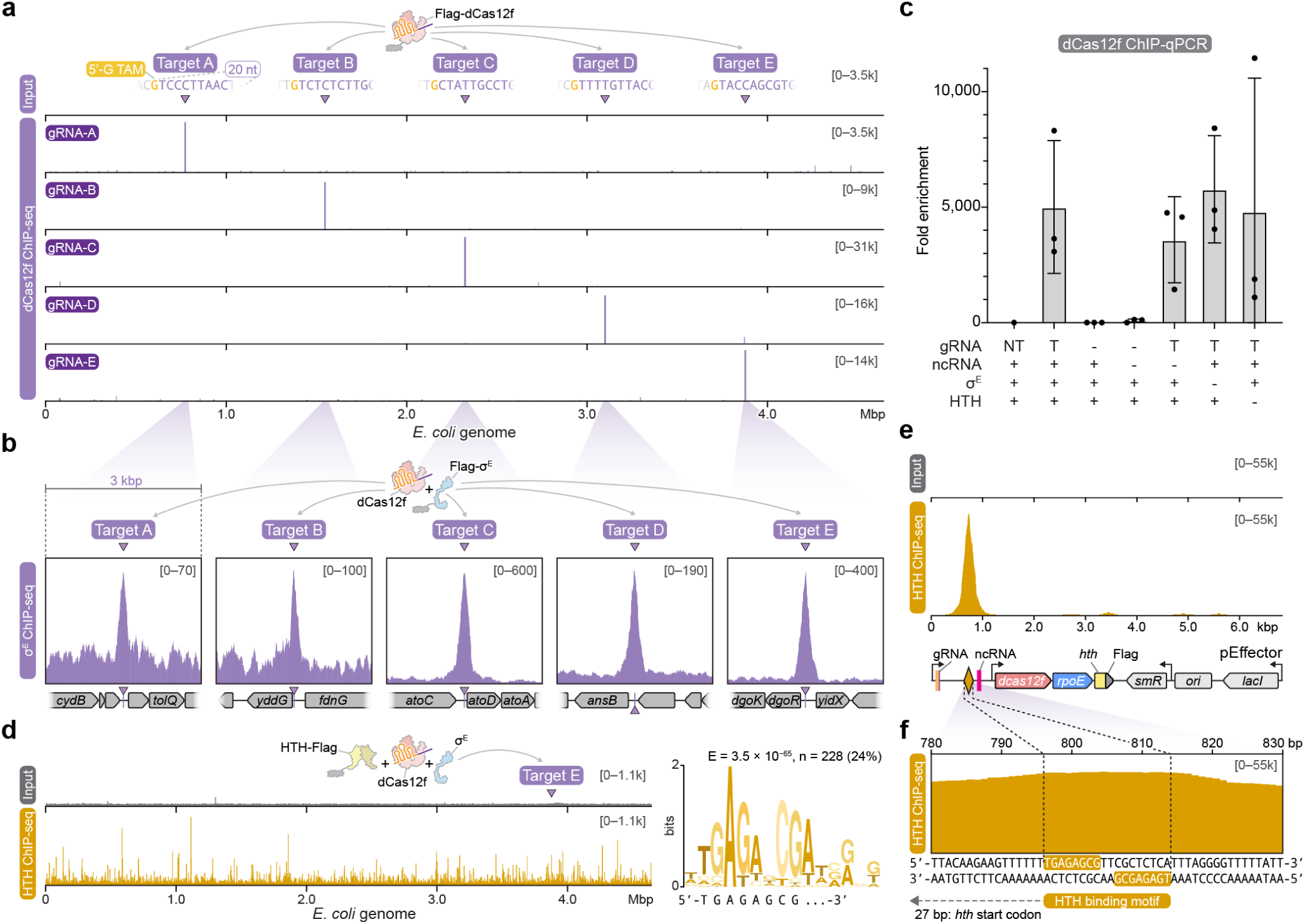
RNA-guided dCas12f recruits σ^E^, but not HTH, to genomic target sites. **a,** Genome-wide representation of ChIP-seq data for *Fta* Flag-dCas12f targeting one of five *E. coli* genomic target sites (A–E, marked with purple triangle) flanked by a 5′-G TAM, alongside the input control (top). Coverage is shown as counts per million (CPM), normalized to the highest peak in the targeting sample. **b,** Magnified view of ChIP-seq data for targets A–E in **a**, but from experiments expressing Flag-σ^E^ and untagged dCas12f, demonstrating that σ^E^ is efficiently recruited to RNA-guided DNA target sites. Genomic regions are annotated below each graph. **c,** Bar graph showing ChIP-qPCR data from experiments in which Flag-dCas12f was tested alongside the indicated conditions. Data are shown as a fold enrichment, calculated from ΔΔC_q_. T, targeting gRNA; NT, non-targeting gRNA; data are mean ± s.d. for *n* = 3 biologically independent samples. **d,** Genome-wide representation of ChIP-seq data (left) for HTH-Flag co-expressed with σ^E^, dCas12f, and a target E-specific gRNA, alongside the input control (top left); coverage is shown as in **a**. Binding events were analyzed by MEME-ChIP (right), which revealed a strongly conserved consensus motif corresponding to the putative native HTH binding site. E, E-value significance; n, number of peaks contributing to the motif. **e,** pEffector plasmid-wide representation of ChIP-seq data from **d**. Coverage is shown as in **a**, and plasmid annotations are shown below the graph. **f**, Magnified view of the peak center from **e**, shown above the underlying DNA sequence. Comparison to the motif from MEME analysis in **c** reveals the likely HTH binding motif comprising two perfectly inverted repeats (yellow boxes, bottom).

Although phylogenetic information disfavored the hypothesis that HTH co-evolved with dCas12f and σ^E^ to form a larger complex, we nevertheless performed experiments to directly immunoprecipitate HTH. The resulting ChIP-seq data revealed widespread binding across the *E. coli* genome, in stark contrast to the binding profiles of dCas12f and σ^E^ (**Fig. 4d**). Subsequent analyses revealed a conserved DNA binding motif that matched a highly enriched sequence found immediately upstream of the *hth* coding sequence, which was cloned into the expression plasmid (**Fig. 4d–f**). This region encompasses inverted repeats that align with the motif, are signature constituents of HTH binding sites^41,49^, and are situated 27 bp from the *hth* start codon, strongly implicating HTH-DNA binding in gene repression. A broader analysis revealed inverted repeats within 50 bp of most *dcas12f*-associated *hth* genes (**Extended Data Fig. 7b**), suggesting that this auto-regulation mode is common in *rpoE-dcas12f* loci.

Collectively, these results demonstrate direct σ^E^ recruitment to genomic sites bound by dCas12f, independent of HTH. To test whether such interactions drive transcription, we next developed a sensitive reporter assay to investigate RNA-guided gene expression.

### RNA-guided transcription by dCas12f and σ^E^

We used our dCas12f-σ^E^ effector plasmid in a reporter assay, wherein we cloned the native target upstream of a red fluorescent protein gene on pTarget (**Fig. 5a**), anticipating that dCas12f-gRNA complexes would recruit σ^E^ to DNA and drive transcription and red fluorescence. Disappointingly, however, initial experiments resulted in no detectable RFP expression above background levels (**Fig. 5b**). Reasoning that *Fta* σ^E^ might be unable to productively engage *Eco*RNAP, we cloned an additional expression plasmid encoding all four *Fta*RNAP subunits (α, β, β’, ω) and repeated the assay (**Fig. 5a**). Remarkably, co-expression of dCas12f, gRNA, σ^E^, and cognate *Fta*RNAP robustly activated RFP expression (**Fig. 5b**). Control experiments demonstrated the necessity of all four components for gene activation, but not HTH and the *hth*-associated ncRNA (**Fig. 5b**). Furthermore, RNA sequencing confirmed strong upregulation of *mRFP* transcription compared to all other *E. coli* genes (**Extended Data Fig. 8a**), and revealed the emergence of a well-defined, novel transcription start site (TSS) 46 bp downstream of the TAM (**Fig. 5c**). Additional controls demonstrated that RFP fluorescence was not the result of transcriptional readthrough, that TAM mutations severely diminished transcriptional activation, and that bulk fluorescence measurements agreed closely with single-cell flow cytometry measurements (**Extended Data Fig. 8b–e, Supplementary Fig. 1**).

**Figure 5.**
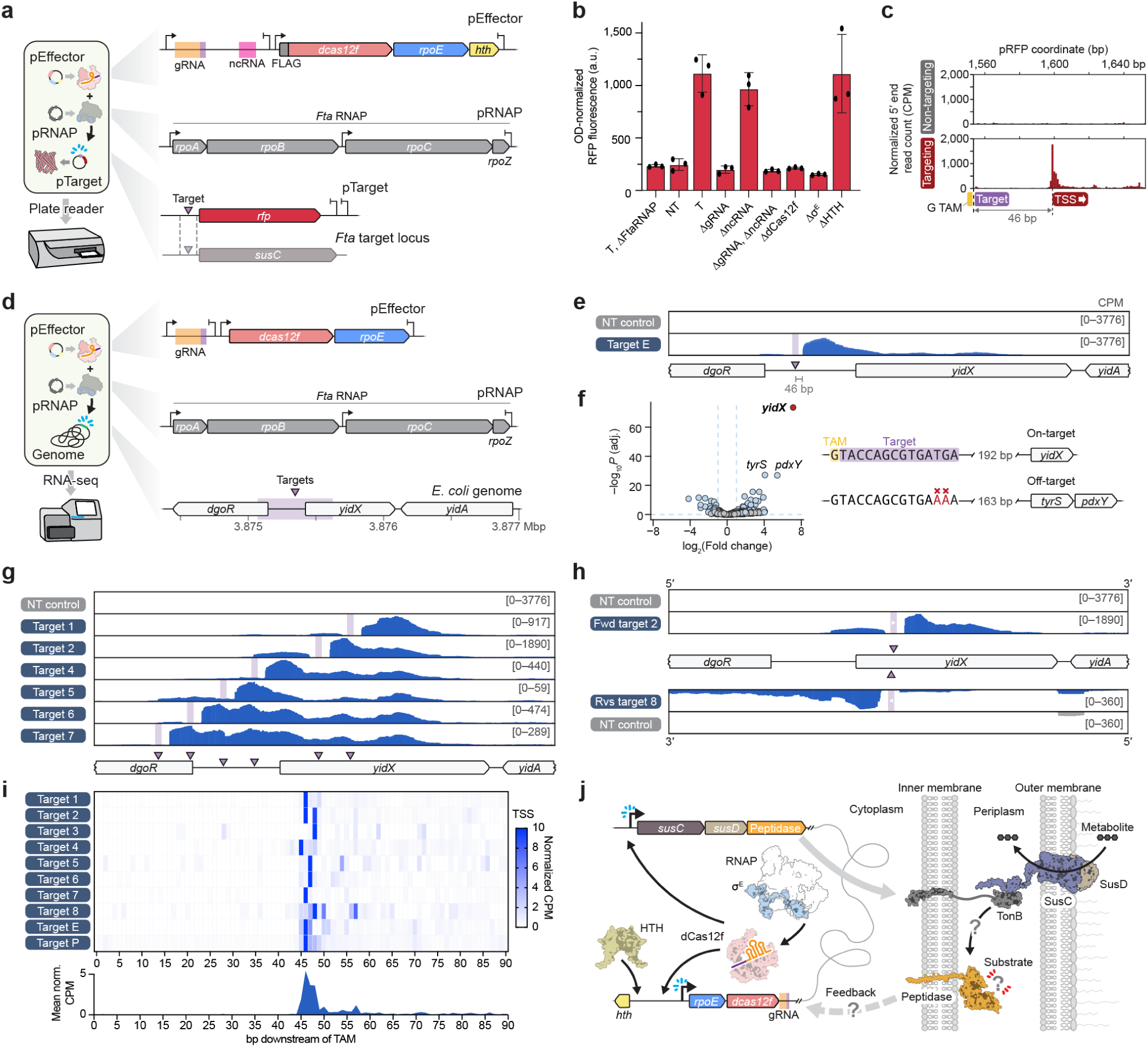
dCas12f and σ^E^ direct programmable, RNA-guided transcription with single-base pair resolution. **a,** Schematic of reporter assay (left) in which protein and RNA components encoded by pEffector and pRNAP drive gene expression of a plasmid-encoded RFP (pTarget), as measured by a fluorescence plate reader. **b,** OD-normalized RFP fluorescence for the indicated experimental conditions using the assay in **a**. T, targeting gRNA; NT, non-targeting gRNA. Data are shown as mean ± s.d. for *n* = 3 biologically independent samples. **c,** Magnified view of RNA-seq data mapped to pTarget for targeting (T) and non-targeting (NT) samples in **b**, shown as the 5′ end read counts per million (CPM). Transcription initiation is tightly confined to a TSS positioned 46 bp downstream of the TAM. **d,** Schematic of transcription assay to drive RNA synthesis from gRNA-matching genomic loci, as measured by RNA-seq. gRNAs were designed to target within an untranscribed region (purple box/triangle) near *dgoR* and *yidX* genes. **e,** RNA-seq coverage for four biological replicates targeting the indicated region (purple triangle), with non-targeting (NT) controls shown. **f,** Differential gene expression analysis for the experiments in **e**, demonstrating that *yidX* is the most significantly up-regulated gene; other up-regulated genes (labeled) are encoded downstream of an identifiable off-target site. Wald test P values adjusted for multiple comparisons were calculated using the Benjamini–Hochberg approach. **g**, RNA-seq coverage with tiled gRNAs targeting sites 1–7, shown as in **e**. **h,** RNA-seq coverage for two gRNAs targeting opposite strands within the *yidX* gene body alongside matched non-targeting controls, shown as in **e**. These results demonstrate that dCas12f can drive transcription in either polarity, depending only on the orientation of the gRNA. **i,** Heat map showing the distance distribution between the TAM and TSS for each of 10 tested gRNAs. The aggregated data is presented in a histogram (bottom), demonstrating that the TSS occurs 46 bp downstream of the TAM. **j,** Model for RNA-guided transcription catalyzed by dCas12f and σ^E^ (*rpoE* gene). gRNAs encoded downstream of *dcas12f* serve to auto-regulate *rpoE-dCas12f*expression and drive transcription of transporter proteins like SusCD, in a mechanism that involves cognate protein-protein interactions between σ^E^, RNAP, and the DNA-bound dCas12f-RNA complex. This mechanism likely evolved to respond to, and regulate, metabolite import during growth in dynamically variable environmental conditions.

Flavobacteria, a heterotrophic group within the phylum *Bacteroidetes*, exhibit consensus promoter motifs centered at positions –7 and –33 relative to the TSS^50,51^, as compared to the –10 and –35 promoter motifs characteristic of *E. coli*. Since this architecture would place the –33 motif within the dCas12f binding footprint, we were curious whether the native target site contained promoter motifs recognized directly by *Fta* σ^E^, or whether gRNA-DNA base pairing provided the sole sequence specificity determinants. Consistent with this latter scenario, bioinformatics analysis of all dCas12f gRNA-matching target sites showed an absence of conserved motifs beyond the presumed RNA-DNA heteroduplex site, apart from a weak thymine preference that may facilitate DNA bending (**Extended Data Fig. 8f**). These clues suggested the intriguing possibility that promoter specificity in dCas12f-σ^E^ systems may rely exclusively on RNA-DNA complementarity, with little or no dependence on direct σ^E^-DNA sequence recognition. To test this, we systematically mutated the region downstream of the guide-target match (**Extended Data Fig. 8g**) and then repeated reporter assays. Strikingly, RFP activation remained largely unchanged with these sequence perturbations, including mutation 8 that scrambled the entire sequence downstream of the target site (**Extended Data Fig. 8h**). These results indicate that sequence-specific σ^E^-DNA interactions are unlikely to be involved in transcription initiation by dCas12f-σ^E^ systems. Structural data from a parallel study suggests that dCas12f-associated σ^E^ engages DNA through non-canonical interactions, explaining reduced requirements for upstream promoter motifs compared to typical σ factors^52^.

Given the strong stimulation of RNA-guided transcription by *Fta*RNAP, we were curious to repeat Flag-σ^E^ ChIP-seq assays in its presence. *Fta*RNAP inclusion substantially increased levels of σ^E^ enrichment at dCas12f-bound target sites, and the sequencing coverage extended well beyond the RNA-matching target site (**Extended Data Fig. 8i**). This result underscores the critical interactions between σ^E^ and its cognate RNAP within the transcription initiation complex^52^, while further reflecting their likely co-migration during the initiation and elongation phases of transcription, prior to σ^E^ being released from the RNAP complex^53^.

Finally, we incrementally reduced RNA-DNA complementarity in 2-bp steps to determine the optimal and minimal guide length required for efficient RNA-guided transcription (**Extended Data Fig. 8j**). We observed strong fluorescence levels for all lengths >10 nt, and a 14-nt guide elicited the strongest RFP expression (**Extended Data Fig. 8k**), in excellent agreement with the *Fta* guide length determined by RIP-seq and RNA-seq (**Fig. 2c,e,h**).

### Programmable RNA-guided gene activation

Given the absence of promoter requirements other than an RNA-DNA match, we hypothesized that gRNAs could be designed to initiate gene expression at arbitrary loci with single nucleotide TSS precision. We focused on a transcriptionally silent intergenic site in *E. coli* where ChIP-seq revealed strong dCas12f and σ^E^ binding (**Fig. 4a,b**), and then assessed RNA-guided transcription by RNA-seq (**Fig. 5d**). Remarkably, our results again revealed precise transcription initiation 46 bp downstream of the TAM, dependent only on a targeting gRNA and 5’-G TAM (**Fig. 5e**). Transcriptome-wide analyses confirmed the high specificity of gene activation (**Fig. 6c**), and RT-qPCR corroborated an approximately 140-fold increase in *yidX*-derived transcripts compared to non-targeting controls (**Fig. 5f, Extended Data Fig. 9a**). We detected significant upregulation of two additional genes — *tyrS* and *pdxY* — but closer inspection revealed a likely off-target site upstream of both genes that exhibited strong similarity to the guide sequence (**Fig. 5f, Extended Data Fig. 9b**). These observations highlight the potency of RNA-guided transcription, the ability to activate multiple genes across operons, and the potential for off-target activity.

Next, building off our hypothesis that RNA-guided transcription does not require fixed promoter motifs, we envisioned that transcription start sites could be predictably shifted within any desired region of interest, simply by redesigning the guide. We generated a panel of seven gRNAs tiled across the *dgoRyidX* locus, with the sole criteria involving a 5’-G TAM and 20 bp of RNA-DNA complementarity. The resulting RNA-seq data strikingly revealed a concomitant shift in the TSS for each tested gRNA (**Fig. 5g, Extended Data Fig. 9c**), with only Target 3 exhibiting intractability, likely due to competing transcriptional regulators binding within this region (**Extended Data Fig. 9d**). RNA-guided transcription strictly required a 5’-G TAM, as activity was lost with gRNAs shifted sequentially by 1 bp to sample each potential TAM at Target 6 (**Extended Data Fig. 9e**). Notably, RNA-guided transcription functioned robustly even when targeting within the *dgoR* and *yidX* gene bodies, where pre-existing promoter motifs would be absent, and we were easily able to drive antisense transcription by designing gRNAs to target the opposite DNA strand (**Fig. 5h**).

To explore the mechanism of RNA-guided transcription, we systematically analyzed TSS distributions across all of the tested gRNAs. Transcription initiation consistently occurred within a narrow window 45–48 bp downstream of the TAM (**Fig. 5i**), suggesting a tight physical coupling between formation of the dCas12f-mediated R-loop and the transcription bubble, as was confirmed by cryo-EM structures of the RNA-guided transcription initiation complex^52^; the presence of some TSS variability indicates that additional sequence determinants may cause genomic context-specific patterns. RNA-guided transcription also consistently produced a slight accumulation of positive-sense RNA-seq reads upstream of the DNA target site (**Fig. 5e,g,h**), which we hypothesize results from occasional loading of *Fta*RNAP onto a nearby chromosomal site in 3D upstream of DNA-bound dCas12f, leading to low-level transcription up until RNAP-dCas12f collisions occur.

Altogether, our results demonstrate that dCas12f-σ^E^ systems drive RNA-guided transcription at sites that can be easily reprogrammed, without any requirement for pre-existing promoter motifs beyond a 5’-G TAM, leading to targeted gene expression of any user-defined locus of interest.

## DISCUSSION

In their foundational work on the *lac* repressor, Jacob and Monod originally theorized that transcription factors (TFs) might be RNAs that could control gene expression through base-pairing interactions^54^. Since then, however, extensive studies have established the prevailing dogma in the field, which primarily ascribes gene regulatory function to protein-based TFs^55^, particularly in the context of transcription initiation and repression. Our work^18^ now demonstrates that diverse gene expression pathways in bacteria — both repressive and activating — leverage guide RNAs to specify genomic targets through RNA–DNA base-pairing, rather than exclusively relying on protein-DNA recognition (**Extended Data Fig. 10**). Regulatory RNAs have previously been shown to modulate gene expression post-transcriptionally by influencing mRNA secondary structure^56^ or stability^57^, but programmable, RNA-guided control of transcription itself represents a novel category of gene regulation. Our finding that dCas12f-σ^E^ systems drive RNA-guided transcription further defines a unique mechanism of transcription initiation, which we speculate could enable more rapid gene evolution via the *de novo* transcription of previously non-coding regions of the bacterial genome.

From our analyses thus far, transmembrane transport operons are frequent native targets of dCas12f-σ^E^ (**Fig. 3d**). These *susCD*-containing loci are typically regulated by a proximally encoded ECF σ factor and its corresponding anti-σ factor, to fine-tune iron uptake, peptide transport, or carbohydrate import as part of polysaccharide utilization loci (PUL)^42,58,59^. Operonically linked nutrient-metabolizing hydrolases often reflect the specific substrate recognized by SusCD transporters^60,61^, but for *Fta* dCas12f-σ^E^, the *sus-CD*-adjacent gene is a peptidase (**Fig. 3d**). PULs can be further regulated by two-component systems (TCS)^62^, which we also identified as common RNA-guided targets (**Fig. 3d**). We therefore hypothesize that dCas12f-σ^E^ systems emerged to regulate peptide, carbohydrate, or iron acquisition pathways, in which metabolite sensing triggers expression of dCas12f-σ^E^ loci, thereby leading to upregulation of genes involved in mediating substrate uptake (**Fig. 5j**). Future efforts will be necessary to identify the specific metabolite(s) affected by dCas12f-σ^E^ regulation, the physiological conditions that induce their native expression, and how the HTH protein and its associated ncRNA are involved. Addressing these questions remains a challenge since Bacteroidetes genomes harbor dozens of PULs, each responsible for the import and utilization of a distinct glycan^63^. We also identified frequent examples of dCas12f-σ^E^ systems that likely stimulate their own expression (**Fig. 3d**), similar to autoregulatory mechanisms for non-dCas12f ECF σ factors^64^, suggesting a positive feedback loop for amplifying transcription signals.

A fundamental question persists: what advantages do dCas12f-σ^E^ systems provide over existing σ factor groups, beyond introducing regulatory flexibility? We hypothesize that their promoter-independent behavior would allow a small set of target genes to possess their own privatized σ^E^, without risking promoter cross-recognition by other σ factors. This property may be physiologically advantageous by tightly regulating energetically costly pathways like transmembrane transport that requires dedicated channel proteins and metabolic enzymes only needed in the presence of select metabolites^65^. Controlling transcription via an RNA-guided mechanism would also facilitate multiplexing and rapid diversification rates, given the relative ease with which specificity can be altered through co-varying mutations.

It is remarkable that TnpB — a transposon-encoded RNA-guided nuclease that was itself exapted during the emergence of RNA-guided transcriptional repressors^18^ — became domesticated by host adaptive immune systems in the form of CRISPR-Cas12f^6,17^, before being exapted yet again during the emergence of RNA-guided transcriptional activators (**Fig. 1b**). In the course of this evolutionary trajectory, the transposon-encoded single-guide ωRNA^10,46^ presumably split into a dual-guide CRISPR RNA and tracrRNA^44^, before collapsing again into the chimeric single gRNAs used by dCas12f. Further evidence for this sequential transformation is found in the characteristic dimeric architecture^40,66^ of dCas12f bound to the σ^E^-RNAP holoenzyme within the full R-loop transcription bubble, reported in an accompanying article^52^. This cryo-EM structure elegantly corroborates our observation of transcription initiation occurring 45–48 bp downstream of the TAM, while further revealing unique interactions between dCas12f and the extended CTD in dCas12f-associated σ^E^ homologs (**Fig. 1f,g**). We identified related homologs that unexpectedly share this domain but are instead associated with restriction endonuclease (RE)-like genes (**Extended Data Fig. 1c**), suggesting that transcription initiation pathways may have co-opted inactivated nucleases from both adaptive and innate immune systems to provide novel modes of DNA specificity.

Numerous CRISPR-based technologies exist to programmably inhibit or activate gene expression, respectively referred to as CRISPRi and CRISPRa^4,5,25^. Strategies developed in mammalian cells rely on fusions of dCas9 to repressor or activator domains like KRAB^26^ or VP64^5^, which function by adding repressive epigenetic marks or remodelling chromatin to facilitate RNA polymerase recruitment^4,5^. In bacteria, promoter occlusion by dCas9 functions robustly for CRISPRi, but CRISPRa methods have been hampered by intrinsic constraints in the ability to selectively recruit RNAP to pre-existing promoter mo-tifs^29^, limiting applications in challenging biological contexts where promoter architectures are unknown. The dCas12f-σ^E^ systems described here circumvent these restrictions by directly coupling RNA-guided DNA recognition to *de novo* transcription initiation, independent of promoters or host transcriptional machinery. We envision that this promoter-agnostic mechanism could enable precise and flexible control of gene expression from virtually any genomic locus of interest that meets minimal TAM requirements, offering new opportunities to activate silent gene clusters, explore functional genomics in non-model bacteria, and rewire regulatory networks with unprecedented resolution. The ability to sensitively define custom transcription start sites may further expand the use of this system for synthetic biology and metabolic engineering applications.

More broadly, our work adds to a growing body of evidence that RNA-guided proteins — once thought to function primarily in DNA cleavage during antiviral defense — have been repeatedly exapted to serve diverse roles in cell biology. From transposon proliferation to adaptive immunity, and now to transcriptional repression and activation, these systems illustrate the remarkable evolutionary plasticity of RNA-guided mechanisms and highlight their untapped potential for genome control.

## METHODS

### Candidate system choice and phylogenetic analyses of σ^E^-dCas12f systems

An initial set of σ^E^-dCas12f systems was identified via a BLASTp^67^ search (-evalue 1e-50 -qcov_hsp_perc 80 -max_hsps 50) of the NR database with a dCas12f homolog (WP_076354695.1) observed during manual analyses of Cas12 loci. The resulting 499 Cas12f-like proteins were then aligned to the structurally characterized *Cn*Cas12f homolog (WP_120361969.1, PDB: 8HR5), and the dCas12f homolog used to query the NR database, confirming that all 499 homologs from the initial search were predicted to have mutations in their RuvC nuclease active sites. These sequences were then used as queries in a subsequent BLASTp^67^ search of the NR database, which identified 161 additional homologs. This joint dataset was de-duplicated, resulting in 660 unique dCas12f homologs, which were merged with σ^E^-associated dCas12f homologs identified in a recent bioinformatic survey of TnpB/Cas12 proteins^17^. Homologs encoded in a genomic contig that precluded extraction of a dCas12f locus at least 30 kbp in length were then removed from this dataset of 869 unique dCas12f proteins. The final dataset of 707 unique dCas12f proteins were aligned with MAFFT^68^ (LINSI option) and a phylogenetic tree was built with FastTree^69^ (-wag -gamma). Sixteen dCas12f homologs were then chosen to sample the phylogenetic diversity of these systems.

To construct a phylogenetic tree of dCas12f proteins and their most closely related nuclease-active relatives, known σ^E^-associated dCas12f proteins were plotted on a phylogenetic tree from a recently published bioinformatic survey of Cas12/TnpB^17^ proteins, and a clade consisting of 99 tips was extracted from this tree. This subtree was made up of Cas12f and dCas12f representative proteins that had been clustered at 50% amino acid identity. To expand this tree, we fetched 1,487 homologs represented by the branches of this tree, aligned them with MAFFT^68^ (LINSI option), and built a new tree with FastTree^69^ (-wag -gamma). RuvC active site intactness and domain composition were annotated for each homolog via comparison to a homolog in the tree that is nearly identical (99.8% aa identity) to the structurally characterized *Un*Cas12f homolog (PDB: 7L48). CRISPR associations were extracted from the published metadata accompanying the previous dCas12f bioinformatic survey by Altae-Tran *et al*.^17^ Associations for *rpoE* and HTH genes were annotated via BLASTp^67^ searches of ORFs occurring upstream of dCas12f, queried with σ^E^ and HTH protein sequences from the 16 systems selected for experimental characterization.

To obtain a set of σ^E^ sequences for phylogenetic analysis, we conducted a BLASTp^67^ search of a local copy of the NCBI non-redundant protein database (-evalue 0.01 -max_target_seqs 10000), queried with the *Fta* σ^E^ sequence (RIV51358.1). The resulting 2,709 sequences were then clustered with the easy-linclust option in MMseqs^70^ at a threshold of 50% amino acid identity and 80% alignment coverage. Next, cluster representatives (n = 872) were aligned with MAFFT^68^ (LINSI option), the alignment was trimmed of columns composed of >50% gaps with Trimal^71^, and a phylogenetic tree was built from the trimmed MSA with FastTree^69^ (-wag -gamma). This tree was used to guide the selection of 247 cluster representatives that formed a monophyletic clade with dCas12f-associated σ^E^s. The 823 members of clusters these sequences represented were then retrieved, aligned (MAFFT^68^ – LINSI), and used to build a tree (FastTree^69^ -wag -gamma). Genomic assemblies encoding each *rpoE* homolog were fetched with the rentrez^72^ package in R, and *rpoE* loci (comprising 20 kbp of genomic sequence) were annotated with Eggnog^73^ mapper to assess the genetic context of each homolog. dCas12f associations were additionally verified using a custom profile hidden Markov model built from dCas12f homologs identified in the analyses described above.

### Cloning and strain generation

All proteins, strains, and plasmids used in this study are described in **Supplementary Tables 4-6**. Initially, genes encoding the σ^E^, dCas12f, and HTH protein were synthesized by Genscript and cloned into pCDFDuet-1 vectors (pEffector) along with intergenic (non-coding) regions flanking dCas12f and σ^E^-coding genes using PfoI and Bsu36I restriction sites (**Fig. 2a**). Three systems encoded two distinct HTH proteins. For those, both HTH-coding genes were cloned onto the same plasmid. A 3×Flag epitope tag was appended to σ^E^, the N-terminus of dCas12f, or the C-terminus of HTH. Expression of these proteins was driven by a constitutive J23105 promoter. Transcription of non-coding regions was driven by a separate constitutive J23119 promoter. The *F. taeanensis* (GCA_003584105.1) RNA polymerase subunits encoded by *rpoA*, *rpoB*, *rpoC* and *rpoZ* were cloned onto a pACYCDuet vector (p*Fta*RNAP) driven by two T7 promoters (**Fig. 5a**). Codon optimization was performed for all genes. Subsequent cloning used a variety of methods, including inverse (around-the-horn) PCR, Gibson assembly, restriction-digestion ligation, and ligation of hybridized oligonucleotides. Q5 DNA polymerase was used for all PCR steps. Plasmids were cloned, grown in NEB Turbo cells, purified with Qiagen miniprep kits, and then verified by Sanger Sequencing (Genewiz).

### RNA immunoprecipitation and sequencing (RIP)

RIP-seq sample preparation was generally performed as described in our previous studies^18,74^ with differences in culturing. Individual *E. coli* K-12 MG1655 (sSL0810) colonies harboring the pEffector plasmid, with either dCas12f or HTH carrying a 3×Flag tag, were inoculated into 25 ml of LB media supplemented with spectinomycin (100 μg ml^-1^). Liquid cultures were grown at 37 °C for approximately 16 h. A volume equivalent to 20 ml of OD_600 nm_ = 0.5 was aliquoted. Cells were harvested by centrifugation at 4,000 g for 5 min, followed by a wash with 1 ml of cold TBS buffer. The supernatant was removed, and the pellet was flash-frozen in liquid nitrogen and stored at −80°C.

For immunoprecipitation, monoclonal anti-Flag M2 antibodies produced in mouse (Sigma-Aldrich, F1804) were conjugated to Dynabeads Protein G (Thermo Fisher Scientific) magnetic beads. For each sample, 60 μl of magnetic beads were washed three times in IP lysis buffer (20 mM Tris-HCl, pH 7.5 at 25°C, 150 mM KCl, 1 mM MgCl_2_, 0.2% Triton X-100) and resuspended in 1 ml IP lysis buffer. Then, 20 μl anti-Flag antibody was added and incubated under rotation for >3 h at 4 °C. Antibody-bead-mixtures were washed two times and resuspended in 60 μl × n samples of IP lysis buffer.

Cell pellets stored at −80°C were thawed on ice and resuspended in 1.2 ml IP lysis buffer supplemented with 1× cOmplete Protease Inhibitor Cocktail (Roche) and 1 μl SUPERase•In RNase Inhibitor (Thermo Fisher Scientific). Cells were lysed by sonication using a 1/8″ sonicator probe with the following settings: 1.5 min total sonication time (2 s on, 5s off), 20% amplitude. Lysates were centrifuged at 21,000 g for 15 min at 4 °C and the supernatant was transferred to a new tube. 10 μl of samples were set aside as non-immunoprecipitated input controls and stored at −80 °C. 60 μl of conjugated antibody-beads slurry was added to the remainder of the supernatant and rotated overnight at 4 °C. After incubation, samples were washed three times with 1 ml of cold IP wash buffer (20 mM Tris-HCl, pH 7.5 at 25°C, 150 mM KCl, 1 mM MgCl_2_) on a magnetic rack. To elute immunoprecipitated RNA, magnetic beads were resuspended in 1 ml TRIzol (Thermo Fisher Scientific) and incubated at RT for 5 min. The supernatant containing eluted RNA was transferred to a new tube and 200 μl of chloroform was added, followed by vigorous mixing. The samples were incubated at RT for 3 min and centrifuged for 15 min at 12,000 g at 4 °C. Then, the RNA Clean & Concentrator-5 kit (Zymo Research) was used to isolate RNA from the upper aqueous layer. RNA was eluted in 15 μl nuclease-free water. RNA from input samples was isolated alongside the RIP samples using TRIzol followed by column purification. Purified RNA was stored at −80 °C prior to next-generation sequencing library preparation.

For RIP-seq library preparation, RNA from input and RIP samples was fragmented by random hydrolysis of 7 μl RNA, 6 μl water, and 2 μl NEBuffer 2 at 92 °C for 2 min. Samples were then treated with 2 μl TURBO DNase (Thermo Fisher Scientific) and 2 μl RppH (NEB) in the presence of 1 μl SUPERase•In RNase Inhibitor for 30 min at 37°C. Next, samples were treated with 1 μl T4 PNK (NEB) in 1× T4 DNA ligase buffer (NEB) for 30 min at 37°C. RNA was column purified using the Zymo RNA Clean and Concentrator-5 kit and eluted in 10.5 μl nuclease-free water. RNA concentrations were determined using the DeNovix RNA Assay Kit. Sequencing libraries were prepared using the NEBNext Small RNA Library Prep Kit. Oligonucleotide and barcode primer sequences used for library generation are listed in **Supplementary Table 7**. Illumina libraries were sequenced in 75×75 and 150×150 paired-end mode on the Illumina NextSeq 500 and Element Biosciences Aviti Cloudbreak Freestyle platforms.

### RIP-seq analyses

Sequencing analyses were generally performed as described previously^74^. Raw and processed sequencing files are described in **Supplementary Table 8**. RIP-seq and input datasets were processed using cutadapt^75^ (v4.2) to trim Illumina adapter sequences, remove low-quality ends, and eliminate reads shorter than 15 bp. Read mapping was performed using a combined reference genome file containing the *E. coli* K-12 MG1655 genome (NC_000913.3) and relevant plasmid sequences using bwamem2^76^ (v2.2.1) with default parameters. Alignments were sorted and indexed using SAMtools^77^ (v1.17). deepTools2^78^ (v3.5.4) bamCoverage was used to generate counts per million (CPM)-normalized coverage tracks with a bin size of 1, with combined top and bottom strand alignments, scaled according to sequencing depth. RIP-seq coverage tracks were visualized in IGV^79^ (v2.14.1). Transcriptome-wide analyses of RIP-seq enrichment at annotated features was performed using featureCounts^80^ (v2.0.2) with -s 1 for strandedness, and further processed using DESeq2^81^ to calculate fold-change and FDR (using the Benjamini-Hochberg approach) between input and RIP sample for each annotated feature. Visualizations were generated using ggplot2.

### Chromatin immunoprecipitation (ChIP)

Sample generation for ChIP was generally performed as previously described^82^, with changes in culturing and sonication conditions. Individual *E. coli* K-12 MG1655 (sSL0810) colonies harboring the pEffector plasmid, with either dCas12f, σ^E^ or HTH carrying a 3×Flag tag, were inoculated into 25 ml of LB media supplemented with spectinomycin (100 μg ml^-1^). For one sample, p*Fta*RNAP was co-transformed with pEffector (**Extended Data Fig. 8i**) and LB media was supplemented with additional chloramphenicol (25 μg ml^-1^). Liquid cultures were grown at 37 °C for approximately 16 h, before fresh LB was added to a final volume of 40 ml. Cells were fixed promptly by addition of 1 ml of 37% (v/v) formaldehyde (Thermo Fisher Scientific) followed by vigorous vortexing and crosslinking continued for 20 min at RT while gently shaking. Crosslinking was quenched by addition of 4.6 ml of 2.5 M glycine, followed by 10 min incubation with gentle shaking. To pellet cells, samples were centrifuged at 4,000 g for 8 min at 4 °C. The next steps were performed on ice. The supernatant was discarded and pellets were resuspended in 40 ml TBS buffer (20 mM Tris-HCl pH 7.5, 0.15 M NaCl). Cells equivalent to 40 ml of OD_600 nm_ = 0.6 were aliquoted into a new tube. Cells were pelleted again by centrifugation and transferred into an Eppendorf tube, flash-frozen in liquid nitrogen and stored at −80 °C or kept on ice prior to the subsequent steps.

Immunoprecipitation was performed as follows: bovine serum albumin (GoldBio) was dissolved in PBS buffer (Gibco) for a final concentration of 5 mg/ml BSA. 25 µl of Dynabeads Protein G (Thermo Fisher) slurry (hereafter referred to as ‘beads’ or ‘magnetic beads’) were prepared for each sample. At the maximum, 250 µl of the initial slurry were processed in a single tube. All bead washes were performed at RT using a magnetic rack to retain the beads. First the initial supernatant was removed. Then, beads were washed four times using 1 ml of BSA solution. Beads were resuspended in 25 µl × n samples of BSA solution, followed by addition of 4 µl × n samples of monoclonal anti-Flag M2 antibody produced in mouse (Sigma-Aldrich, F1804). Antibody and magnetic beads were conjugated for >3 h at 4 °C while rotating. In the meantime, crosslinked cell pellets were thawed on ice and resuspended in FA lysis buffer 150 (50 mM HEPES-KOH pH 7.5, 0.1% (w/v) sodium deoxycholate, 0.1% (w/v) SDS, 1 mM EDTA, 1% (v/v) Triton X-100, 150 mM NaCl), supplemented with protease inhibitor cocktail (Sigma-Aldrich) and transferred to a 1 ml milliTUBE AFA Fiber (Covaris). Next, samples were sonicated on a LE220 Focused-ultrasonicator (Covaris) using a 24 place milliTUBE rack (Covaris) with the following SonoLab 7.3 settings: min. temp. 4 °C, set point 6 °C, max. temp. 8 °C, peak power 420, duty factor 30, cycles/bursts 200, 17.5 min sonication time. Sonicated samples were centrifuged at 20,000 g and 4 °C for 20 min. The supernatant was transferred into a new tube and kept on ice. 10 µl of the supernatant was then set aside as a non-immunoprecipitated input control, flash-frozen in liquid nitrogen, and stored at −80 °C. After >3 h of conjugating antibodies to magnetic beads, the slurry was washed four times with 1 ml of BSA solution, as described earlier. Then, magnetic beads were resuspended in 30 µl × n samples of FA lysis buffer 150 with protease inhibitor, and 31 µl of resuspended antibody-conjugated beads were added to each sonicated cell lysate sample, followed by overnight incubation at 4 °C while rotating to immunoprecipitate the Flag-tagged proteins. Then next day, beads were washed using 1 ml of buffers, as follows: two washes with FA lysis buffer 150, one wash with FA lysis buffer 500 (50 mM HEPES-KOH pH 7.5, 0.1% (w/v) sodium deoxycholate, 0.1% (w/v) SDS, 1 mM EDTA, 1% (v/v) Triton X-100, 500 mM NaCl), one wash with ChIP wash buffer (10 mM Tris-HCl pH 8.0, 250 mM LiCl, 0.5% (w/v) sodium deoxycholate, 0.1% (w/v) SDS, 1 mM EDTA, 1% (v/v) Triton X-100, 500 mM NaCl), and two washes with TE Buffer 10/1 (10 mM Tris-HCl pH 8.0, 1 mM EDTA). Finally, immunoprecipitated protein-DNA complexes were eluted by addition 200 µl of freshly made ChIP elution buffer (1% (w/v) SDS, 0.1 M NaHCO_3_) and incubation for 1.25 h at 65 °C with vortexing every 15 min. In the meantime, input samples were thawed and 190 µl of ChIP elution buffer was added, followed by the addition of 10 µl of 5 M NaCl. After the 1.25 h incubation, magnetic beads were discarded and the supernatant was transferred to a new tube, and 9.75 µl of 5 M NaCl were added. ChIP and input samples were incubated at 65°C overnight to de-crosslink protein-DNA complexes. Then, RNA was degraded by addition of 1 µl of 10 mg/ml RNase A (Thermo Fisher) and incubation for 1 h at 37 °C, followed by protein degradation by addition of 2.8 µl of 20 mg/ml Proteinase K (Fisher Scientific) and 1 h incubation at 55 °C. 1 ml of buffer PB (QIAGEN) was added to all ChIP and input samples, followed by column purification and elution in 40 ul TE Buffer 10/0.1 (10 mM Tris-HCl pH 8.0, 0.1 mM EDTA).

For ChIP-qPCR, pairs of genomic qPCR primers (**Supplementary Table 7**) were designed for the *yidX* target site (tSL0679) and a genomic reference locus (*rssA*). 1 µl of ChIP or input sample was diluted with 14 µl of TE buffer 10/0.1. Parallel qPCR reactions were performed using both genomic target and reference locus primer pairs and the fold change between ChIP and input samples was calculated using the ΔΔ*C*_q_ method: *A* = *C*_q_(immunoprecipitated sample at target locus) − *C*_q_(immunoprecipitated sample at reference locus); *B* = *C*_q_ (input sample at target locus) − *C*_q_ (input sample at reference locus); ΔΔ*C* = 2^−(*A*−*B*)^.

For ChIP-seq, libraries were generated using the NEBNext Ultra II DNA Library Prep Kit for Illumina (NEB). Initial DNA concentrations were quantified using the DeNovix dsDNA Ultra High Sensitivity Kit. DNA amounts were standardized across ChIP and input samples prior to library preparation. Adapters were ligated and 12 to 15 cycles of PCR amplification were performed to add Illumina barcodes (**Supplementary Table 7**). Then, two-sided size selection using AMPure XP beads (Beckman Coulter) was performed to purify DNA fragments of around 450 bp length, as described previously^82^. For sample pooling, final DNA concentrations were quantified using the DeNovix dsDNA High Sensitivity Kit. Illumina libraries were sequenced in 75×75 and 150×150 paired-end mode on the Illumina NextSeq 500 and Element Biosciences Aviti Cloudbreak Freestyle platforms.

### ChIP-seq analyses

Sequencing analyses were performed as described previously^82^. Raw and processed sequencing files are described in **Supplementary Table 8**. ChIP-seq reads were trimmed using fastp^83^ (v0.23.2) and mapped to a custom *E. coli* K-12 MG1655 reference genome using Bowtie2^84^ (v2.2.5) with default parameters. A genomic *lacZ*/*lacI* region and flanking sequences partially identical to plasmid-encoded sequences were masked in all alignments (genomic coordinates 366,386-367,588). Mapped read sorting and indexing was performed using SAMtools^77^ (v1.6). Read coverage was normalized by CPM (Counts Per Million) and binned in 1-bp windows using deepTools2^78^ (v3.5.4) bamCoverage. Normalized reads were visualized in IGV (v2.14.1)^79^. Genome-wide views showing maximum enrichment values per 1-kbp window, as described previously^85^, were generated using bedtools^86^ (v2.26.0) and the ‘bedtools makewindows -w 1000 command’. Motif analysis was performed as described in our previous study^82^. ChIP-seq peaks were called on read alignments prior to normalization using MACS3^87^ (v3.0.2), relative to non-immunoprecipitated input control samples. Peak summit coordinates were extracted together with 100 bp sequences flanking on either side. MEME ChIP^88^ (v5.5.7) using default parameters was used to determine protein-bound sequence motifs.

### Whole genome sequencing (WGS)

One *M. rigui* and five *F. taeanensis* strains were obtained from the China General Microbiological Culture Collection Center (CGMCC) and the Marine Culture Collection of China (MCCC), respectively (**Supplementary Table 5**). Freeze-dried *M. rigui* cells were resuspended in liquid PCA media (5 g tryptone/l, 2.5 g yeast extract/l, 1 g glucose/l) and grown in 5 ml of PCA media at 25 °C for 4 days without antibiotic selection. A volume equivalent to 10 ml of OD_600 nm_ = 1.0 was aliquoted for harvesting. Cells were pelleted by centrifugation at 4,000 g for 5 min, followed by a wash with 1 ml of PBS buffer and another 5 min centrifugation step. Cell pellets were resuspended in 0.5 ml of 1X DNA/ RNA shield buffer (Zymo Research). WGS and gene annotation was performed by Plasmidsaurus.

Freeze-dried *F. taeanensis* strains were resuspended in Difco Marine Broth 2216 and streaked out onto Difco Marine Broth 2216 agar plates. Agar plates were incubated at 28 °C for 3 days. Next, 20 ml of liquid Marine Broth was inoculated from a single colony and grown for another 3 days at 28 °C without antibiotic selection. Further sample preparation was performed as described for *M. rigui*.

### Total RNA sequencing (RNA-seq)

Culturing of *F. taeanensis* was performed as described for WGS. For *E. coli* RNA-seq, BL21(DE3) cells with a genomically encoded *sfGFP* harboring a pTarget plasmid and the *Fta*RNAP plasmid, an empty vector or no plasmid, were transformed with a pEffector plasmid encoding the native *Fta* gRNA or gRNAs with re-programmed guide portions to target sites in the *E. coli* genome. *E. coli* cells were grown in 25 ml of LB media supplemented with antibiotics (100 μg ml^-1^ spectinomycin; 100 μg ml^-1^ carbenicillin; 25 μg ml^-1^ chloramphenicol) and 0.01 mM IPTG for *Fta*RNAP induction. After incubating for 16-20 h at 28 °C, cells equivalent to 2 ml of OD_600 nm_ = 0.5 were harvested by centrifugation at 4,000 g and 4 °C for 5 min. Cell pellets were washed with 1 ml of cold TBS buffer. The supernatant was removed, and the pellet was flash-frozen in liquid nitrogen and stored at −80°C.

Total RNA-seq sample preparation was performed as described previously^74^. Cell pellets stored at −80°C were thawed on ice, resuspended in 1 ml TRIzol (Thermo Fisher Scientific), and incubated at RT for 5 min. Then, 200 μl of chloroform was added, followed by vigorous mixing. The samples were incubated at RT for 3 min and centrifuged for 15 min at 12,000 g at 4 °C. Then, the RNA Clean & Concentrator-5 kit (Zymo Research) was used to isolate RNA from the upper aqueous layer. RNA was eluted in 15 μl nuclease-free water. Purified RNA was stored at −80 °C prior to next-generation sequencing library preparation.

For RNA-seq library preparation, RNA was fragmented by random hydrolysis of 7 μl RNA, 6 μl water, and 2 μl NEBuffer 2 at 92 °C for 2 min. Samples were then treated with 2 μl TURBO DNase (Thermo Fisher Scientific) and 2 μl RppH (NEB) in the presence of 1 μl SUPERase•In RNase Inhibitor for 30 min at 37°C. Next, samples were treated with 1 μl T4 PNK (NEB) in 1× T4 DNA ligase buffer (NEB) for 30 min at 37°C. RNA was column purified using the Zymo RNA Clean and Concentrator-5 kit and eluted in 10.5 μl nuclease-free water. RNA concentrations were determined using the DeNovix RNA Assay Kit.

Sequencing libraries were prepared using the NEBNext Small RNA Library Prep Kit. Illumina libraries were sequenced in 75×75 and 150×150 paired-end mode on the Illumina NextSeq 500 and Element Biosciences Aviti Cloudbreak Freestyle platforms.

### RNA-seq analyses

Sequencing analyses and visualization were performed as described for RIP-seq analyses, with the exception for read mapping and normalization. Raw and processed sequencing files are described in **Supplementary Table 8**. Read mapping using bwa-mem2^76^ (v2.2.1) required a seed sequence of 25 nt for read mapping (-k 25) to avoid gRNA guide sequence mapping to genomic or plasmid target sites. The following *E. coli* BL21(DE3) reference genome was used for read mapping: CP001509.3. Counts per million (CPM)-normalized coverage tracks were generated using deepTools2^78^ (v3.5.4) bamCoverage with a bin size of 1, with combined top and bottom strand alignments, or separation of top and bottom strand alignments. Coverage was scaled according to sequencing depth. DESeq2^81^ was used to calculate fold-change and FDR (using the Benjamini-Hochberg approach) between NT and *yidX* targeting samples for each annotated feature. Differential expression analyses shown in **Figure 5f** employed four biological replicates, and the genomic coordinates for *yidX* were extended to encompass the 5’-UTR observed in RNA-seq data shown in **Figure 5e**, 161 bp upstream of the *yidX* start codon. Normalized CPM in **Figure 5i** was calculated as follows: normalized CPM = (CPM at coordinate *n*) / ((max. CPM in 90-bp TSS window) / 10). Mean normalized CPM was calculated by taking the mean at every coordinate in the same 90-bp window around the TSS, across the ten RNA-seq samples shown in **Figure 5i**.

### RT-qPCR

Purified RNA (pre-library preparation) samples for RNA-seq were used to perform RT-qPCR. The procedure was generally performed as described previously^74^. RNA was treated with 1 μl of dsDNase (Thermo Fisher Scientific) in 1× dsDNase buffer, with a final reaction volume of 10 μl. Samples were incubated at 37 °C for 2 min, followed by adding 1.1 μl of 100 mM DTT and 5 min incubation at 55 °C to stop the reaction. The iScript cDNA Synthesis Kit (BioRad) was used for reverse transcription. cDNA samples were stored at −20 °C. For qPCR, cDNA samples were diluted 10-fold in nuclease-free water. qPCR was performed using 10 μl reactions containing the following components: 5 μl of SsoAdvanced Universal SYBR Green Supermix (BioRad), 2 μl of 2.5 μM primer pair, 1 μl of nuclease-free water, and 2 μl of 10-fold diluted cDNA. Primer pairs were designed to span the intergenic sequence upstream of the *E. coli yidX* gene (**Supplementary Table 7**). Primers designed for the genomic *rssA* gene served as a reference pair for normalization (**Supplementary Table 7**). Reactions were set up in 384-well PCR plates (BioRad). Measurements were obtained using a CFX384 RealTime PCR Detection System (BioRad). The abundance of RNA expressed from the *yidX* locus was calculated relative to the *rssA* locus using the ΔΔ*C*_q_ method (see ChIP-qPCR).

### RFP fluorescence assay

*E. coli* BL21(DE3) cells with a genomically encoded *sfGFP* harboring a pTarget plasmid in addition to the *Fta*RNAP plasmid, an empty vector or no plasmid, were transformed with a pEffector plasmid encoding the native *Fta* gRNA or gRNAs with re-programmed guide portions to target sites in the *E. coli* genome. pTarget was designed to encode a *mRFP1* gene downstream of 500 bp of the native dCas12f target region from the *F. taeanensis* genome (GCA_003584105.1) followed by a strong synthetic ribosome binding site (RBS). *E. coli* cultures were grown in triplicates by inoculating 5 ml of LB in 24-well plates or 0.5 ml of LB in 96-well plates with single colonies. *Fta*RNAP expression was induced with IPTG (0.01 mM). Cultures were grown at 28 °C for 20-24 h. After incubation, 200 μl of culture was transferred into a 96-well optical bottom plate (Thermo Scientific) for endpoint measurements. The OD_600 nm_ was measured in parallel with mRFP fluorescence signal using a Synergy Neo2 microplate reader (Biotek).

### Flow cytometry analysis

*E. coli* BL21(DE3) cultures for flow cytometry analyses were grown as described for the RFP fluorescence assay. After 20-24 h incubation, cells equivalent to 100 μl of OD_600 nm_ = 1.0 were aliquoted into a conical bottom 96-well plate. Cells were pelleted by centrifugation at 4,000 g for 5 min and resuspended in 100 μl flow cytometry buffer (1× PBS, 1 mM EDTA pH 8.0, 1% (w/v) BSA). Then, cells were pelleted again by centrifugation and resuspended in 200 μl flow cytometry buffer. Resuspended cells were then transferred into a round bottom 96-well plate. Flow cytometry analysis was performed using a Novocyte Penteon instrument. Bacteria were gated on FSC-A and FSC-H first to remove debris, and then on FSC-A and SSC-A to analyze singlets (**Supplementary Fig. 1**). Gating was optimized using GFP-only, RFP-only, GFP+RFP, and no GFP/RFP control samples. Then, mean fluorescence intensity was taken for RFP. Flow cytometry data was analyzed using the NovoExpress software.

### Covariation model building and native target identification

To build a covariance model (CM) of dCas12f gRNA scaffold sequences, scaffolds observed in RIP-seq from seven diverse dCas12f systems (*Fta*, *Lby*, *Cgl*, *Lpa*, *Mri*, *Pum*, *Smi*) were aligned with LocaRNA^89^, then Infernal^90^ was used to build a CM (Scaffold CM) (**Extended Data Fig. 6a**). A separate CM was also built from sequences similar to a ncRNA observed in RNA-sequencing data from sSL4759 (*Fta* strain), which encodes a locus that is closely related to the *Fta* dCas12f locus (95.6% nucleotide identity). The ncRNA sequence from sSL4759 was queried against dCas12f loci, resulting in the identification of 65 similar sequences. These hits were used to build an HMM of ncRNA sequences^91^, which when searched against the same dCas12f resulted in the identification of 140 ncRNA-like sequences. Finally, these sequences were aligned with LocaRNA, and a covariance model (ncRNA-like CM) was built with Infernal (**Extended Data Fig. 6a**).

To identify dCas12f gRNA sequences and their expected targets (**Extended Data Fig. 6b**), a subset of 106 dCas12f encoding genomes were downloaded from NCBI. These genomes were selected because they represented “complete” or “chromosome” level assemblies, and each genome was re-annotated with BAKTA^92^. The Scaffold CM was then used to identify gRNA scaffold sequences, and 14 nucleotide sequences abutting the 3’-end of these scaffolds were extracted as putative gRNA guide sequences, consistent with the length of the *Fta* guide observed in RIP-seq. BLASTn^67^ (-evalue 100 -word_size 7-reward 1 -penalty −3 -gapopen 5 -gapextend 2) was then used to match each guide to potential targets in its originating genome. The pool of potential targets was filtered to remove sequences with one or more mismatches over the theoretical duplex formed by six nucleotides of TAM-proximal gRNA guide and the DNA target sequence (i.e., a seed sequence). Potential targets were also filtered to remove BLASTn hits that overlapped with dCas12f guide coordinates, that did not occur in an intergenic region, or that were upstream of a gene improperly oriented — relative to the dCas12f target — to enable sense-stranded transcription of the gene. The genomic contexts of the remaining 626 potential dCas12f targets were then analyzed by extracting 500 nucleotides of sequence 5’ of the putative target, and 10 kilobases of sequence 3’ of the target. The ncRNA-like CM was used to annotate putative ncRNAs in these loci, and loci were manually scored for similarity, based on the identity of genes immediately downstream of the putative dCas12f gRNA-target duplex.

### *In silico* prediction of HTH binding sites

To identify putative HTH binding sites, intergenic regions comprising less than 1 kbp between the HTH-encoding gene and ncRNAs identified with the ncRNA-like CM described above were extracted from dCas12f loci. An initial nHMMer^91^ search of these 349 loci with the sequence of the *Fta* HTH site (identified via ChIP-seq) did not produce any significant hits outside of the *Fta* locus and genetically identical loci. Therefore, we used the palindrome tool from the EMBOSS^93^ suite to identify inverted repeats in each intergenic region (-minpallen 7 -maxpallen 9 -gaplimit 2 -nummismatches 1). The starting position of palindromes that were similar in length to the *Fta* HTH site were then plotted in R using the ggplot2^94^ package.

### Statistics analyses and figure plotting

All sequencing coverage plots were generated in IGV. Plots showing genome-wide expression and differential expression analyses were generated using custom R scripts. Transcription start site (TSS) plots and heatmaps were generated using a custom python script mapping 5’ or 3’ end coordinates of sequencing reads.

### Data availability

Next-generation sequencing data are available in the National Center for Biotechnology Information (NCBI) Sequence Read Archive (BioProject accession: PRJNA1247282) and the Gene Expression Omnibus (GSE293889). The published genomes used for bioinformatics analyses were obtained from NCBI (**Supplementary Table 4**). Datasets generated and analyzed in the current study are available from the corresponding authors on reasonable request.

### Code availability

Custom scripts used for bioinformatics are available at https://github.com/sternberglab/Hoffmann_et_al_2026

## Supporting information

Supplementary Information Guide

## ACKNOWLEDGMENTS

We thank A.J. Robinson, S. Kang., J.L. Ramirez, and T.M. Smith for laboratory support; E.A. Campbell and S.A. Darst for helpful discussion; Z. Hua for custom scripts; Z. Quan at Fudan University for providing *Fta* strains; the JP Sulzberger Columbia Genome Center for NGS support; and L.F. Landweber for qPCR and gel imager instrument access. S.T. was supported by a Ruth L. Kirchstein Individual Predoc-toral Fellowship (F30AI183830) from the NIH. L.C. was supported by NIH grant R01GM138675 and by the National Science Foundation (NSF) Faculty Early Career Development Program (CAREER) Award 2339799. S.H.S. was supported by NIH grant R01EB031935, NSF Faculty Early Career Development Program (CAREER) Award 2239685, a Pew Biomedical Scholarship, an Irma T. Hirschl Career Scientist Award, the Howard Hughes Medical Institute Investigator Program, and a generous startup package from the Columbia University Irving Medical Center Dean’s Office and the Vagelos Precision Medicine Fund.

## AUTHOR CONTRIBUTIONS

F.T.H. and S.H.S. conceived the project. F.T.H. designed and performed most experiments. T.W. performed most bioinformatics analyses. F.T.H., A.I.P., and J.G.-K. performed cloning and RFP activation assays. R.X. cloned RNAP expression plasmids and contributed to data interpretation. F.T.H. and S.T. analyzed RNA-seq and RIP-seq data. H.C.L. and C.M. performed initial phylogenetics and bioinformatics analyses. G.D.L. and F.T.H. performed flow cytometry assays. L.C. contributed to data interpretation. S.H.S. oversaw the project. F.T.H., T.W., and S.H.S. discussed the data and wrote the manuscript, with input from all authors.

## COMPETING INTERESTS

S.H.S. is a co-founder and scientific advisor to Dahlia Biosciences, a scientific advisor to CrisprBits and Prime Medicine, and an equity holder in Dahlia Biosciences and CrisprBits. S.H.S., F.T.H., and T.W. are inventors on patents related to CRISPR-Cas-like systems and uses thereof. The other authors declare no competing interests.

Correspondence and requests for materials should be addressed to S.H.S.

