## Extended Data Figures 1-10 for "Exapted CRISPR-Cas12f homologs drive RNA-guided transcription"

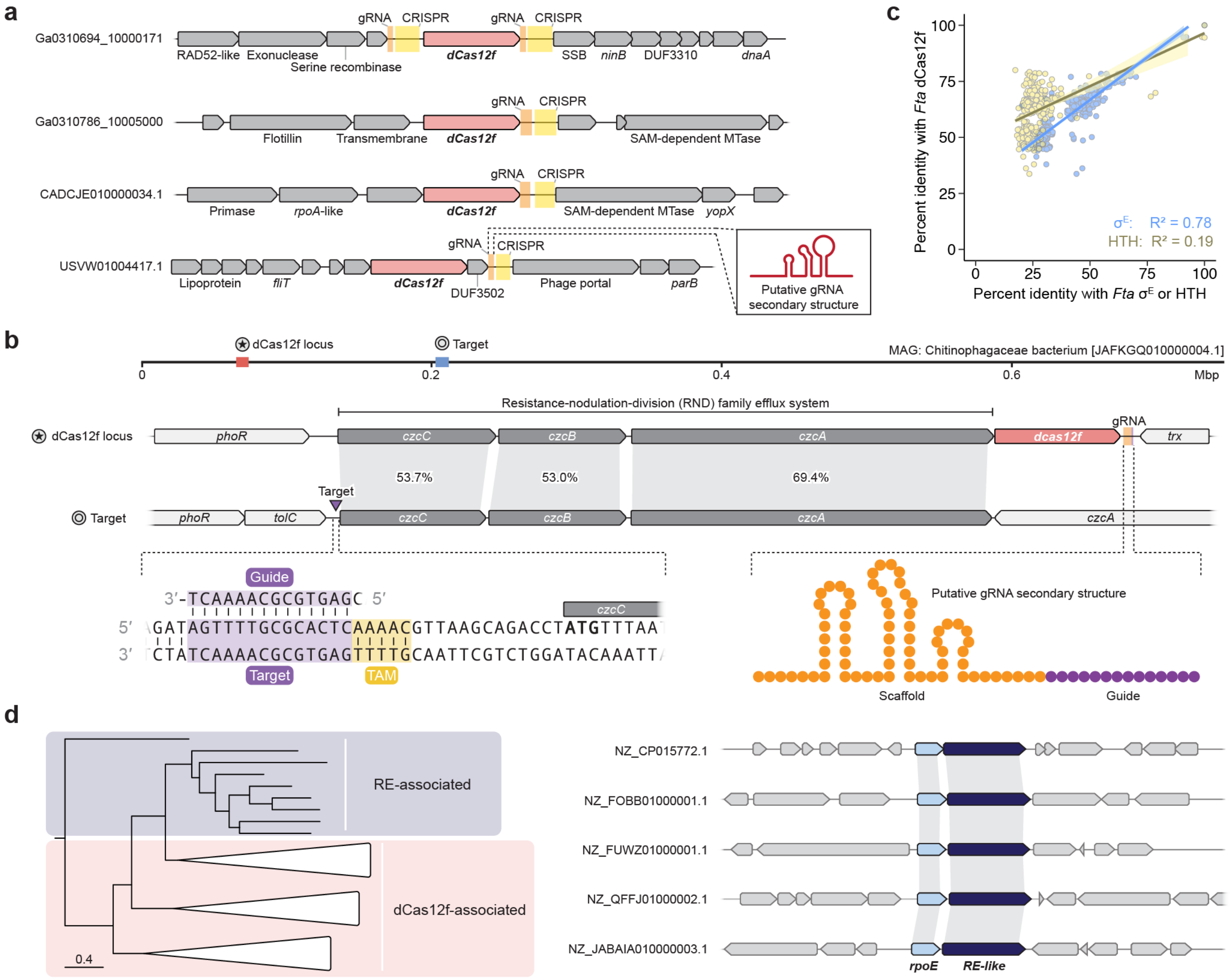


**Extended Data Figure 1 | Evolutionary analysis and genomic context of diverse *dcas12f* and *rpoE* genes. a,** Representative genomic neighborhoods of predicted nuclease-dead Cas12f homologs that are not associated with *rpoE* ($\sigma^E$) genes. CRISPR arrays and putative gRNAs are annotated, as are nearby genes. Putative gRNAs were identified by detecting covariance in the intergenic regions upstream of CRISPR loci, and the predicted secondary structure of a representative example is shown in the inset. **b,** Map of a dCas12f locus and its putative gRNA target upstream of an RND efflux system (middle), in a metagenome assembled genome of a Chitinophagaceae bacterium (top). Putative guide-target duplex and predicted gRNA structures are highlighted (bottom). **c,** Correlation between the percent identity of select dCas12f homologs (y-axis) and either $\sigma^E$ (*rpoE*) or HTH homologs (x-axis) to the *F. taeanensis* homologs. The stronger correlation with $\sigma^E$ ($R^2 = 0.78$) and weaker correlation with HTH ($R^2 = 0.19$) suggest a tighter genetic linkage between *dcas12f* and *rpoE* genes rather than *hth*. **d,** Magnified and simplified view of partial $\sigma^E$ phylogenetic tree from **Fig. 1f** (left), showing homologs containing an additional C-terminal domain (CTD) that are associated with either dCas12f homologs (red) or restriction enzyme (RE)-like homologs (blue). Representative genomic neighborhoods (right) highlight the tight operonic arrangement of *rpoE* and *RE-like* genes, supporting a potential model in which nuclease-dead RE proteins similarly recruit atypical $\sigma^E$ proteins to sites of transcription.

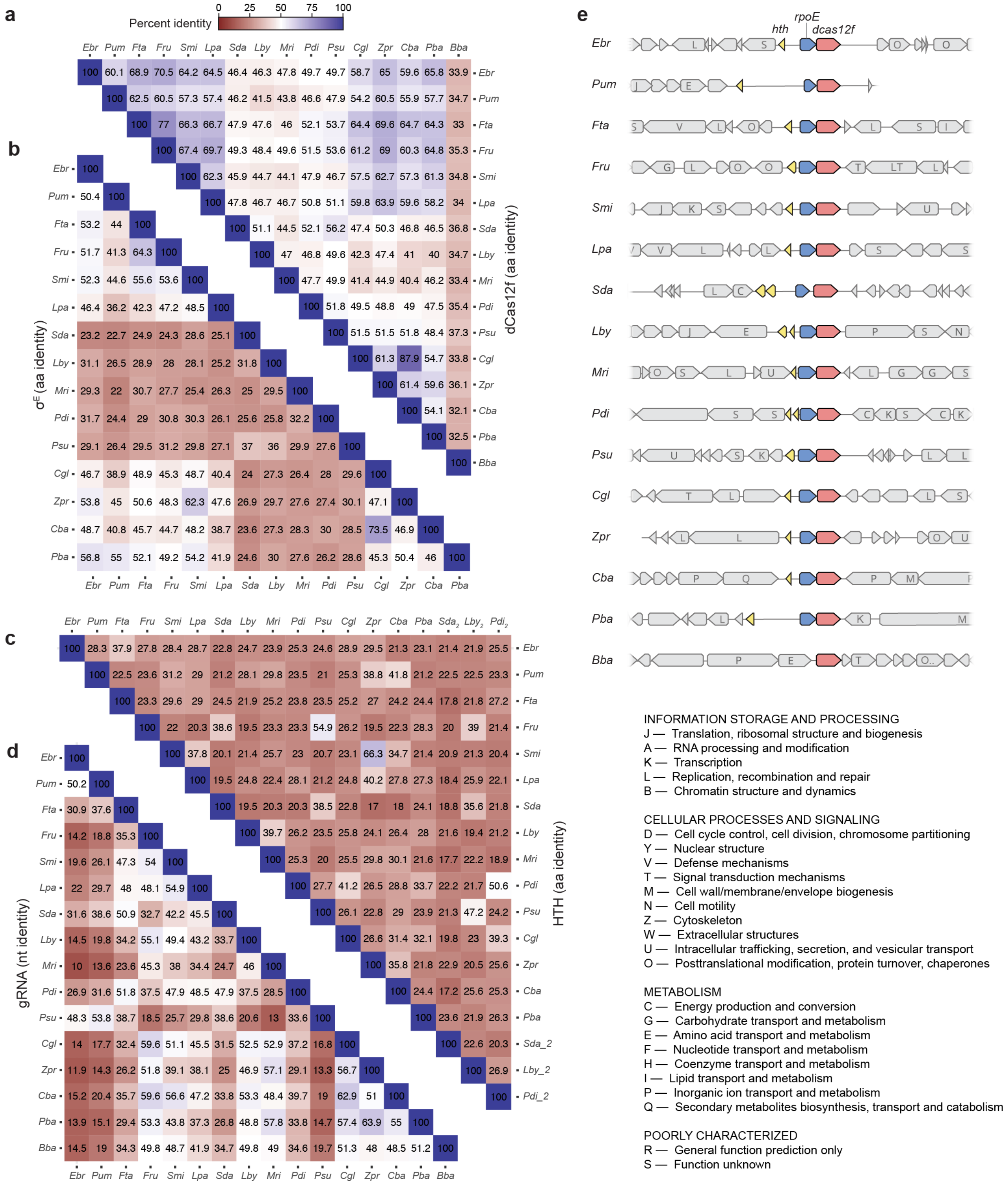


**Extended Data Figure 2 | Pairwise sequence identity matrices for dCas12f, σ$^E$, HTH, and gRNAs. a,** Heatmap of pairwise amino acid sequence identity percentages among dCas12f homologs tested in this study. The matrix is color-coded by sequence identity (see legend inset), and percentages are listed. **b,** Pairwise amino acid sequence identity between tested σ$^E$ homologs, shown as in **a**. The *Bba* locus lacks an *rpoE* gene. **c,** Pairwise amino acid sequence identity between tested HTH homologs, shown as in **a**. Note that *Sda, Lby,* and *Pdi* loci contain two *hth* genes, while this gene is absent in the *Bba* locus. **d,** Pairwise nucleotide sequence identity between tested gRNAs, shown as in **a**. **e,** Annotated genomic loci of all homologous dCas12f-σ$^E$ systems tested in this study, with *hth*, *rpoE*, and *dcas12f* genes labeled and colored in yellow, blue, and red, respectively. Predicted functions for other neighboring genes in grey are indicated by COG (Clusters of Orthologous Groups) letters, as indicated in the legend below.

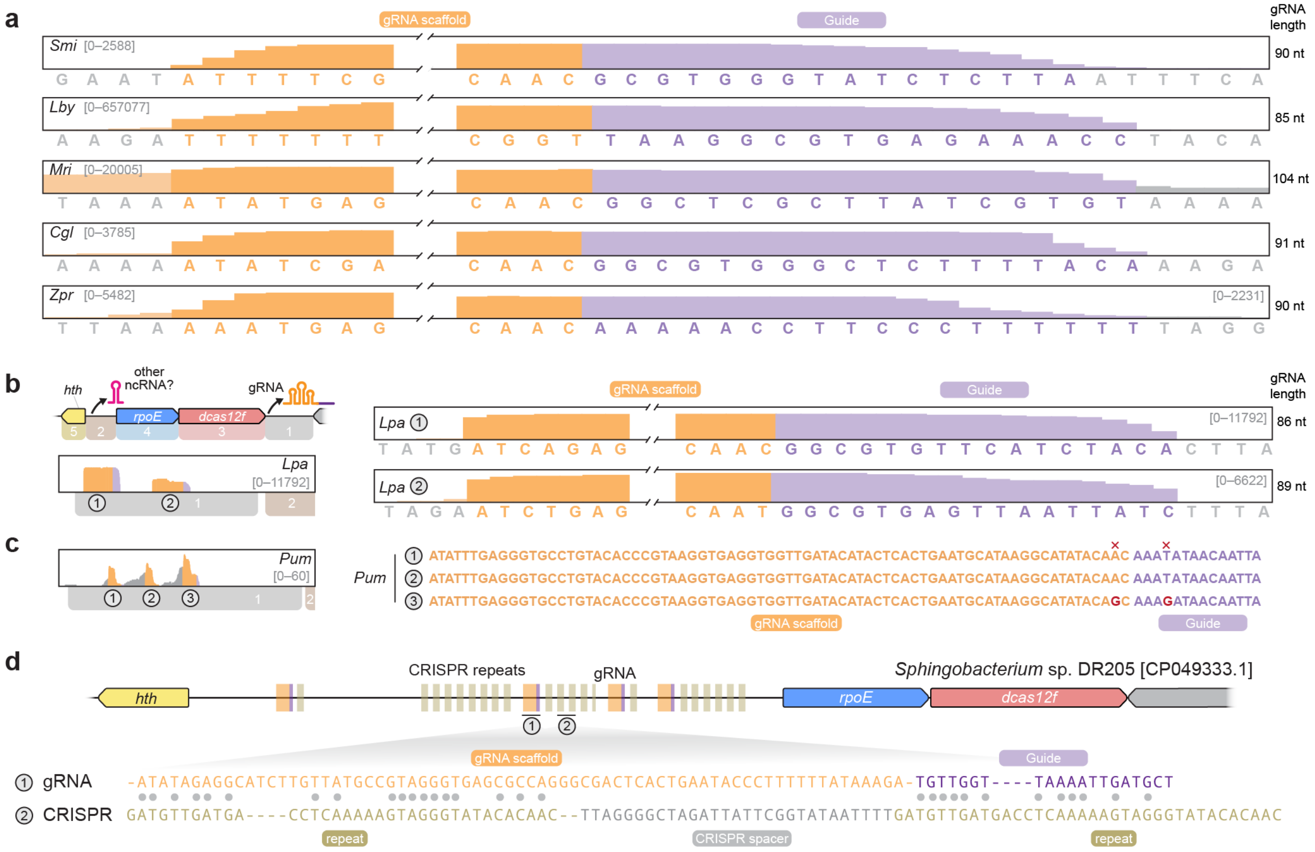


**Extended Data Figure 3 | Additional analysis of dCas12f-associated gRNAs from RIP-seq experiments and comparative genomics analyses. a,** RIP-seq coverage plots for the five indicated dCas12f homologs, revealing a well-defined gRNA with scaffold (orange) and guide (purple) regions. Nucleotides outside the boundary of the presumed full-length gRNA are colored in grey. Total gRNA lengths are noted to the right of each plot. The guide region of *Zunongwangia profunda* is plotted on a separate Y-axis scale, for visual clarity. **b,** Zoomed-out RIP-seq coverage plots for an additional dCas12f ortholog from *Leeuwenhoekiella palythoae* (*Lpa*), showing both a zoomed-out view (left), depicting the same data as shown in **Fig. 2c**, and magnified view as in **a** (right). Region 1 annotated in the operon schematic (top left) encodes multiple tandem gRNAs with similar scaffold and guide sequences. **c,** RIP-seq coverage plots for an additional dCas12f ortholog from *Paenimyroides ummariense* (*Pum*), shown as in **b**, with the magnified view at right comparing the aligned scaffold and guide sequences. The first two gRNA sequences are identical, and bases that differ in the third gRNA are highlighted in red. **d,** Annotated genomic neighborhood of an *rpoE-dcas12f* operon that encodes both predicted full-length gRNAs (orange/purple) and multiple discontinuous CRISPR arrays (repeats in tan). The sequence similarity between the CRISPR repeats and gRNA scaffold (grey circles, bottom) suggests a potential evolutionary emergence of chimeric, dCas12f-associated single-guide RNAs from CRISPR arrays.

**a**

| Sternberg strain ID (sSL) | sSL4575 | sSL4757 | sSL4758 | sSL4759 | sSL4760 | sSL4761 |
|---|---|---|---|---|---|---|
| Strain name | *Mucilaginibacter rigui* | *Flagellimonas taeanensis* | *Flagellimonas taeanensis* | *Flagellimonas taeanensis* | *Flagellimonas taeanensis* | *Flagellimonas taeanensis* |
| Strain bank source | CGMCC (China) | MCCC (China) | MCCC (China) | MCCC (China) | MCCC (China) | MCCC (China) |
| Strain bank ID | 1.13878T | 1K06693 | 1K06752 | 1K06699 | 1K06755 | 1K06720 |
| Growth media | Plate count agar (PCA) | Difco Marine Broth 2216 | Difco Marine Broth 2216 | Difco Marine Broth 2216 | Difco Marine Broth 2216 | Difco Marine Broth 2216 |
| Optimum growth temperature | 25 °C | 28 °C | 28 °C | 28 °C | 28 °C | 28 °C |
| Number of *rpoE-dcas12f* loci | 1 | 0 | 2 | 2 | 0 | 0 |
| Genome accession | GCA_014773275.1 | - | - | - | - | - |

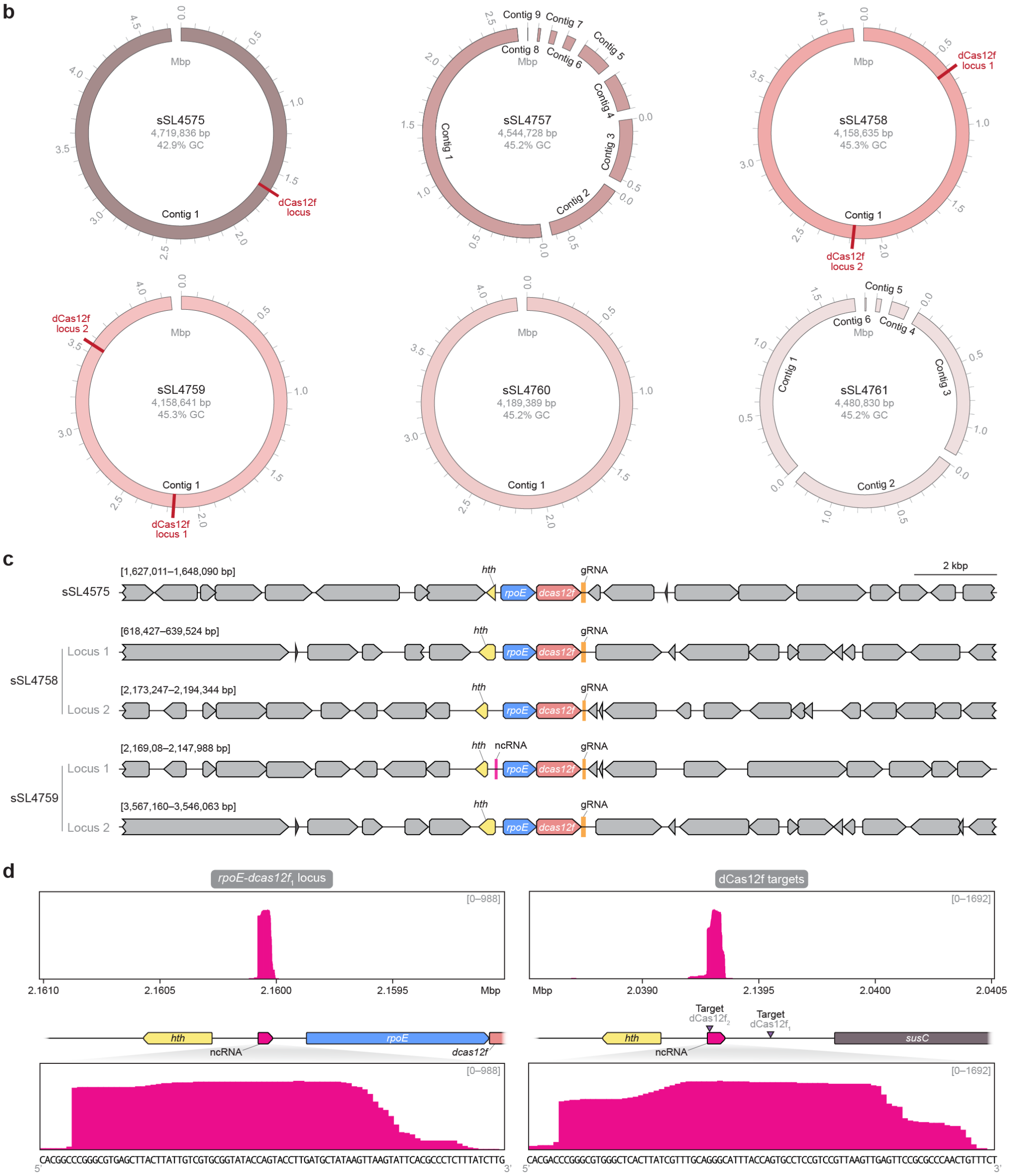


**Extended Data Figure 4 | Culturing, whole genome sequencing, and RNA-seq of *Flagellimonas taeanensis* strains that encode dCas12f-σ$^E$ systems. a,** Summary table of strain information and culturing conditions for five *F. taeanensis* (*Fta*) strains and one *Mucilaginibacter rigui* strain, which were acquired because of the likely presence of *rpoE-dcas12f* loci. The number of loci identified after whole-genome sequencing (WGS) is highlighted. **b,** BioCircos plots of the six strains in **a** after WGS analysis, with the positions of *rpoE-dcas12f* loci highlighted in red. Information regarding the internal strain ID, total genome size, GC content; contigs are denoted in cases where genome assembly was incomplete. **c,** Genomic neighborhoods of *rpoE-dcas12f* loci in the indicated strains from **a**. Genes encoding HTH (*hth*), σ$^E$ (*rpoE*) and dCas12f (*dcas12f*) are shown in yellow, blue, and red, respectively; gRNAs and *hth*-associated ncRNAs are annotated in orange and magenta, respectively. **d,** Magnified RNA-seq coverage plots from *Fta* strain sSL4759 for two distinct loci, highlighting the abundance of reads corresponding to *hth*-associated ncRNAs at both *rpoE-dcas12f* locus 1 (left) and the presumed *susC* target site of dCas12f-associated gRNAs (right). The top panels show a 2-kbp window; the bottom panels zoom in on the ncRNA sequence.

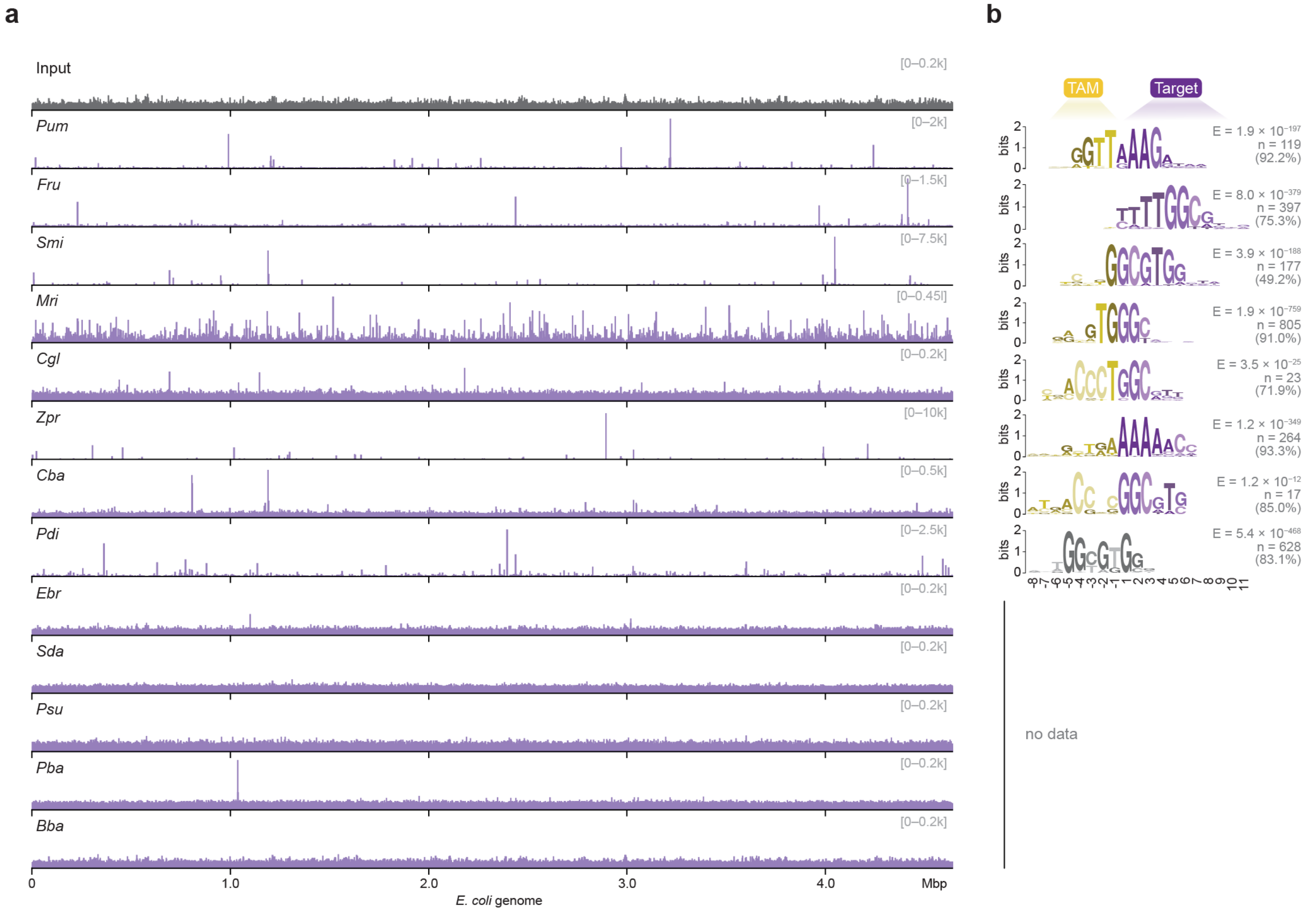


**Extended Data Figure 5 | ChIP-seq experiments reveal TAM and gRNA guide sequences for additional dCas12f homologs. a,** Genome-wide representation of ChIP-seq data for the indicated dCas12f homologs (purple), compared to the input control scaled the same as *Ebr* (top). Coverage is shown as counts per million (CPM), normalized to the highest peak within each sample or to a value of 200, as shown. **b,** Binding events were analyzed by MEME-ChIP, which revealed strongly conserved consensus motifs for eight dCas12f homologs that correspond to the putative target-adjacent motif (TAM) and gRNA-matching target DNA sequence within the seed, for each homolog. E, E-value significance; n, number of peaks contributing to the motif. Percent of total peaks constituent for each motif are shown in parentheses. Motifs could not be confidently determined for the remaining five dCas12f homologs due to a paucity of enriched peaks in heterologous expression experiments.

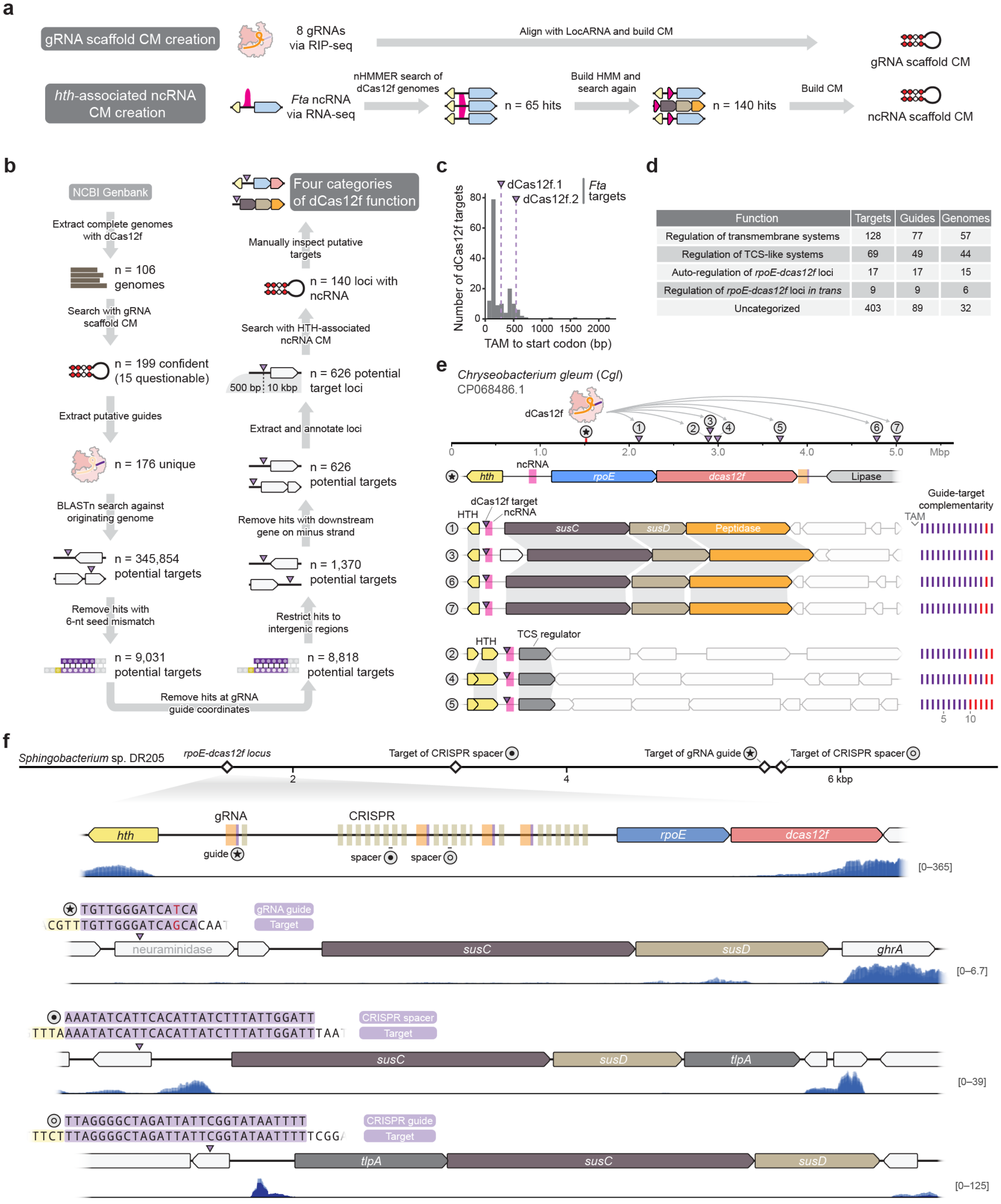


| Function | Targets | Guides | Genomes |
|---|---|---|---|
| Regulation of transmembrane systems | 128 | 77 | 57 |
| Regulation of TCS-like systems | 69 | 49 | 44 |
| Auto-regulation of *rpoE-dcas12f* loci | 17 | 17 | 15 |
| Regulation of *rpoE-dcas12f* loci *in trans* | 9 | 9 | 6 |
| Uncategorized | 403 | 89 | 32 |

**Extended Data Figure 6 | Investigative strategy to uncover putative regulatory functions of dCas12f-σ$^E$ systems. a,** Bioinformatics strategy to determine high-confidence covariance models (CM) for both the dCas12f-associated gRNA (top) and *hth*-associated ncRNA (bottom). **b,** Schematic of bioinformatics workflow to globally identify RNA-guided DNA targets of dCas12f-σ$^E$ systems in sequenced bacterial genomes. After identifying dCas12f-associated gRNAs and extracting guide sequences, putative targets were identified that exhibit perfect complementarity within a 6-nt seed sequence, reside within intergenic regions, and exist upstream of protein-coding genes. Target loci were also analyzed for the presence of predicted *hth*-associated ncRNAs. **c,** Histogram quantifying distances between the TAM of predicted dCas12f target sites and the start codons of associated target genes. 156 bioinformatically predicted dCas12f targets were included in this analysis; distances for Fta dCas12f.1 and dCas12f.2 are highlighted for reference. **d,** Table summarizing the bioinformatics results from **b**, listing the number of predicted DNA targets, gRNA guide sequences, and genomes for each class. Putative RNA-guided DNA targets fall into four functional categories that include regulation of transmembrane transport, regulation of two-component system (TCS)-like systems, auto-regulation of *rpoE-dcas12f* loci, and regulation of *rpoE-dcas12f* loci *in trans*; other predicted targets await further categorization and analysis. Note that some guides have multiple targets within a genome, so a single guide is represented in multiple functional categories and totals do not match totals in **b**. Each guide within a genome is also capable of targeting multiple loci. Thus, some genomes have gRNAs with targets spanning multiple functional classes and totals similarly do match totals in **b**. **e,** Exemplary dCas12f-$\sigma^E$ system from *Chryseobacterium gleum* (*Cgl*), in which the dCas12f-associated gRNA putatively targets and transcriptionally regulates seven distinct genetic loci (1–7). The genome schematic (top) visualizes the approximate genomic location of each locus, and the magnified insets (below) report the position of dCas12f-gRNA targets (purple triangles) relative to nearby genes; note that each of the targets flanks a nearby *hth* gene and overlaps precisely with the predicted position of an *hth*-associated ncRNA (magenta rectangle). The schematics at right depict patterns of gRNA-DNA complementarity at each target site, relative to the TAM. **f,** Exemplary dCas12f-$\sigma^E$ system from *Sphingobacterium* sp. DR205, in which both a single chimeric gRNA guide and two spacers from a vestigial CRISPR array target genomic sites proximal to *susCD* operons. The genome schematic (top) visualizes the approximate location of the *rpoE-dcas12f* locus and putative target sites, and the magnified insets (below) visualize the *rpoE-dcas12f* locus and RNA-guided DNA target sites, alongside corresponding published RNA-seq data for each locus. Three guides/spacers are indicated and labeled (circles), as well as their complementary targets (purple triangles), the predicted guide-target complementarity (purple shading), and the putative TAM (yellow shading).

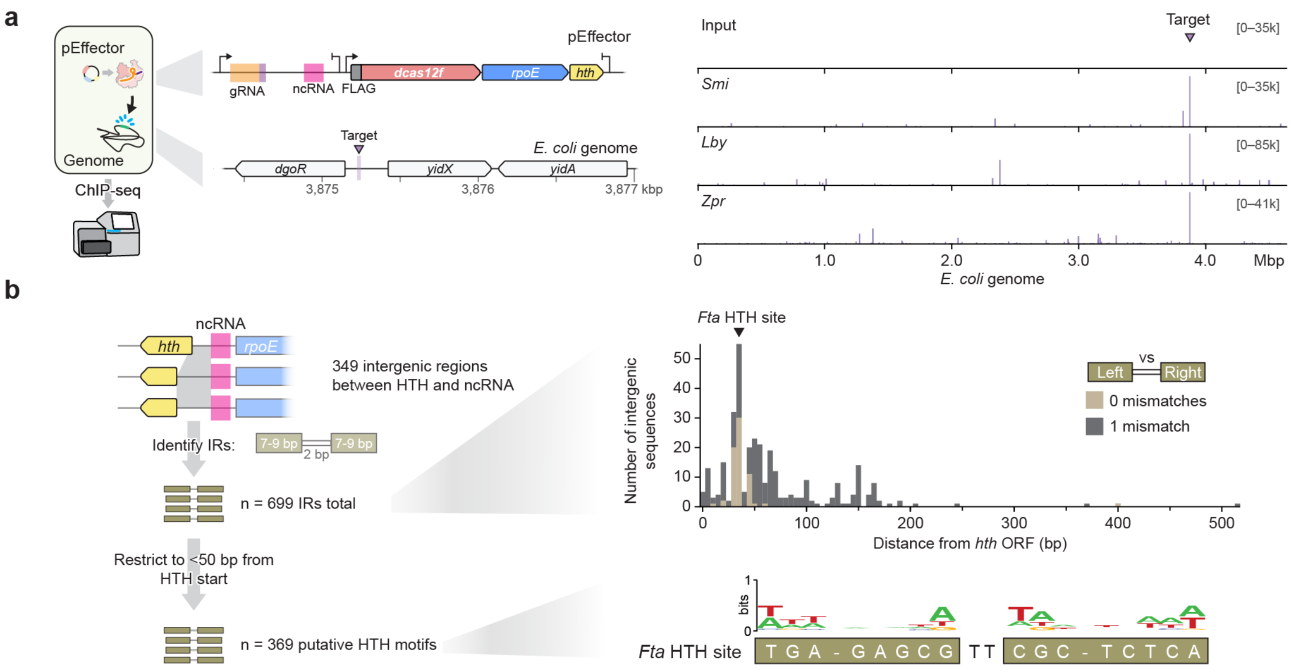


**Extended Data Figure 7 | ChIP-seq experiments and analyses for dCas12f and HTH homologs. a,** Schematic of ChIP-seq assay to study genome-wide binding of dCas12f homologs programmed with gRNAs targeting the indicated site upstream of *yidX* (left), and genome-wide representation of ChIP-seq data alongside the input control (right). Coverage is shown as counts per million (CPM), normalized to the highest peak in the targeting samples. **b,** Bioinformatics strategy to globally discover putative binding sites of HTH proteins (left). Based on ChIP-seq data for *Fta* HTH that revealed a highly enriched binding site with inverted repeats (IRs) upstream of its own open reading frame (ORF), 369 additional potential HTH motifs were identified that similarly exhibit IRs but share little conservation (WebLogo, bottom right), consistent with the broad diversity of *hth* genes encoded in *rpoE-dcas12f* loci. The histogram of IR mismatch (top right) plots the number of mismatches between left and right copies of putative IR substrates of HTH, and their relative distance from the start of the *hth* ORF.

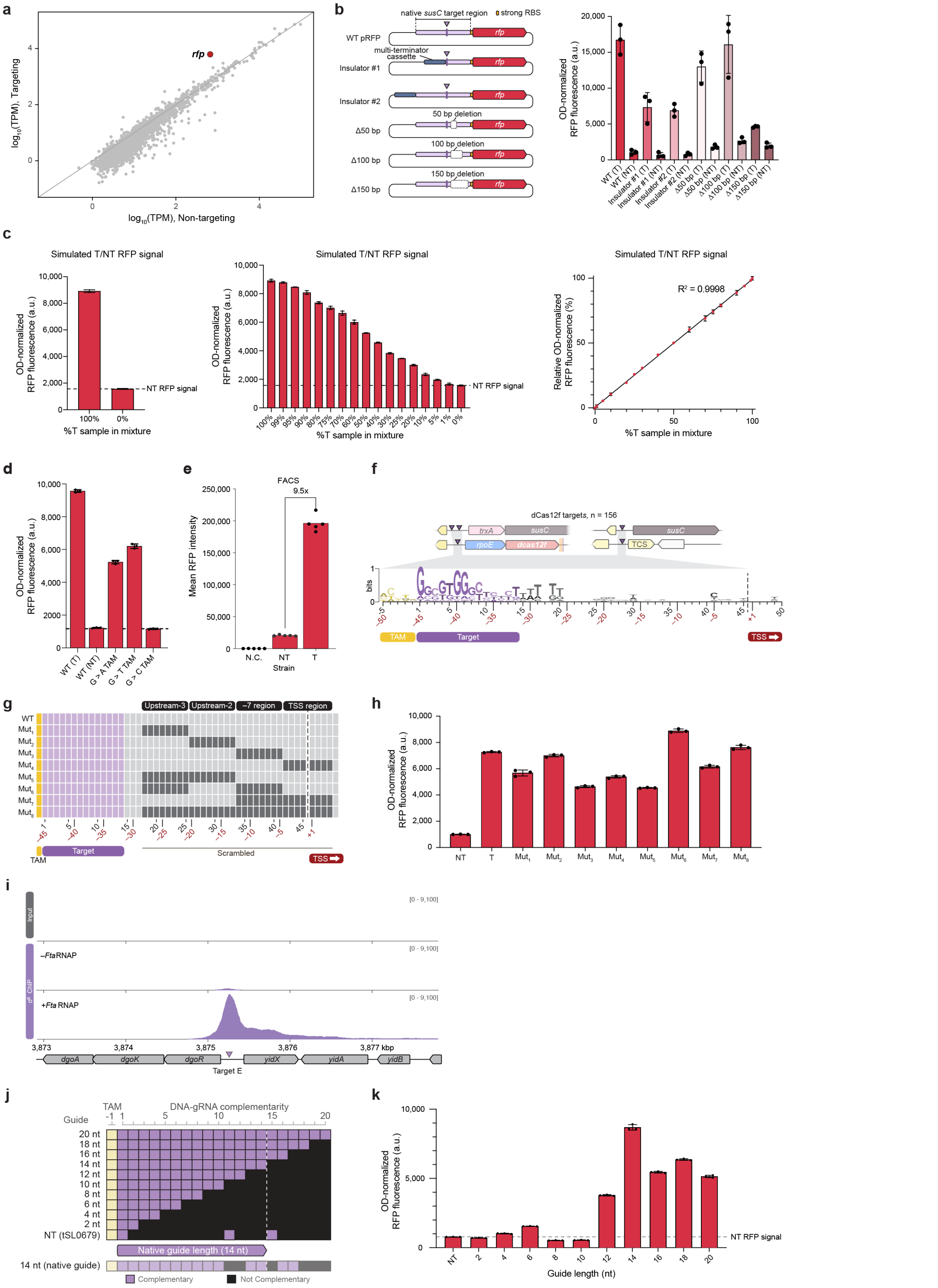


**Extended Data Figure 8 | Optimization and additional analyses relating to RFP fluorescence reporter and ChIP-seq data. a,** Scatter plot of RNA-seq data from mRFP transcriptional reporter assays in **Fig. 5a**, comparing three replicates of targeting (T) or non-targeting (NT) gRNA experiments. TPM, transcripts per million. **b,** Panel of modified mRFP reporter plasmids (left), in which perturbations were made within the native *susC* target region including deletions or multi-terminator cassette insertions, as indicated. The resulting OD-normalized RFP fluorescence measurements for transcriptional reporter assays using these constructs are shown (right), with the starting (WT) construct testing in both targeting (T) and non-targeting (NT) conditions. **c,** OD-normalized RFP fluorescence measurements from control assays to determine signal detection limits. Cell culture samples from targeting (T) and non-targeting (NT) experiments (left) were mixed together to simulate variable ratios of RFP signal (middle), and the resulting regression analysis (right) revealed the expected linear relationship, with excellent sensitivity down to low single-digit percentages. **d,** OD-normalized RFP fluorescence for the indicated reporter DNA constructs, in which the TAM was mutated to each of the alternative nucleotides while maintaining an invariant target and guide sequence. **e,** Flow cytometry analysis for T and NT samples from **b**, alongside a negative control (N.C.) encoding no *mRFP1*. **f,** Consensus WebLogo of dCas12f gRNA-matching target sites from 156 aligned genomic loci, demonstrating conservation within the TAM, the target region, and a short T-rich stretch immediately adjacent to the target — but an absence of substantial conservation in other promoter motifs or the TSS itself. This observation suggests that RNA-guided transcription proceeds largely without fixed DNA sequence requirements. Coordinates are numbered relative to the TAM (top, black text) or to the TSS (bottom, red text). **g**, Panel of mutations in the region between the RNA-matching target site and empirically determined TSS, which were tested in the transcription reporter assay shown in **Fig. 5a**. Nucleotides highlighted in dark grey were mutated; coordinates are numbered relative to the TAM (top, black text) or to the TSS (bottom, red text). **h,** OD-normalized RFP fluorescence for the indicated reporter DNA constructs shown in **g**. T, targeting gRNA with WT (unmutated) intergenic region; NT, non-targeting gRNA control. **i,** Magnified view of ChIP-seq data for Flag-tagged $\sigma^E$ in the absence (-*Fta*RNAP) and presence (+*Fta*RNAP) of the native *Fta*RNAP, alongside an input control (top). Coverage is shown as counts per million (CPM), normalized to the highest peak in the +*Fta*RNAP sample. **j,** Panel of guide sequence truncations in 2-bp increments, which were tested in the transcription reporter assay shown in **Fig. 5a**. The nucleotides highlighted in brown were truncated from the targeting gRNA. **k,** OD-normalized RFP fluorescence for the indicated gRNA constructs shown in **j**, using the transcriptional reporter assay shown in **Fig 5a**. NT, non-targeting gRNA control. Data in **b**, **h,** and **k** are shown as mean ± s.d. for $n = 3$ biologically independent samples. Data in **c, d,** and **e** are shown as mean ± s.d. for $n = 3$, $n = 3$, and for $n = 5$ technical replicates, respectively.

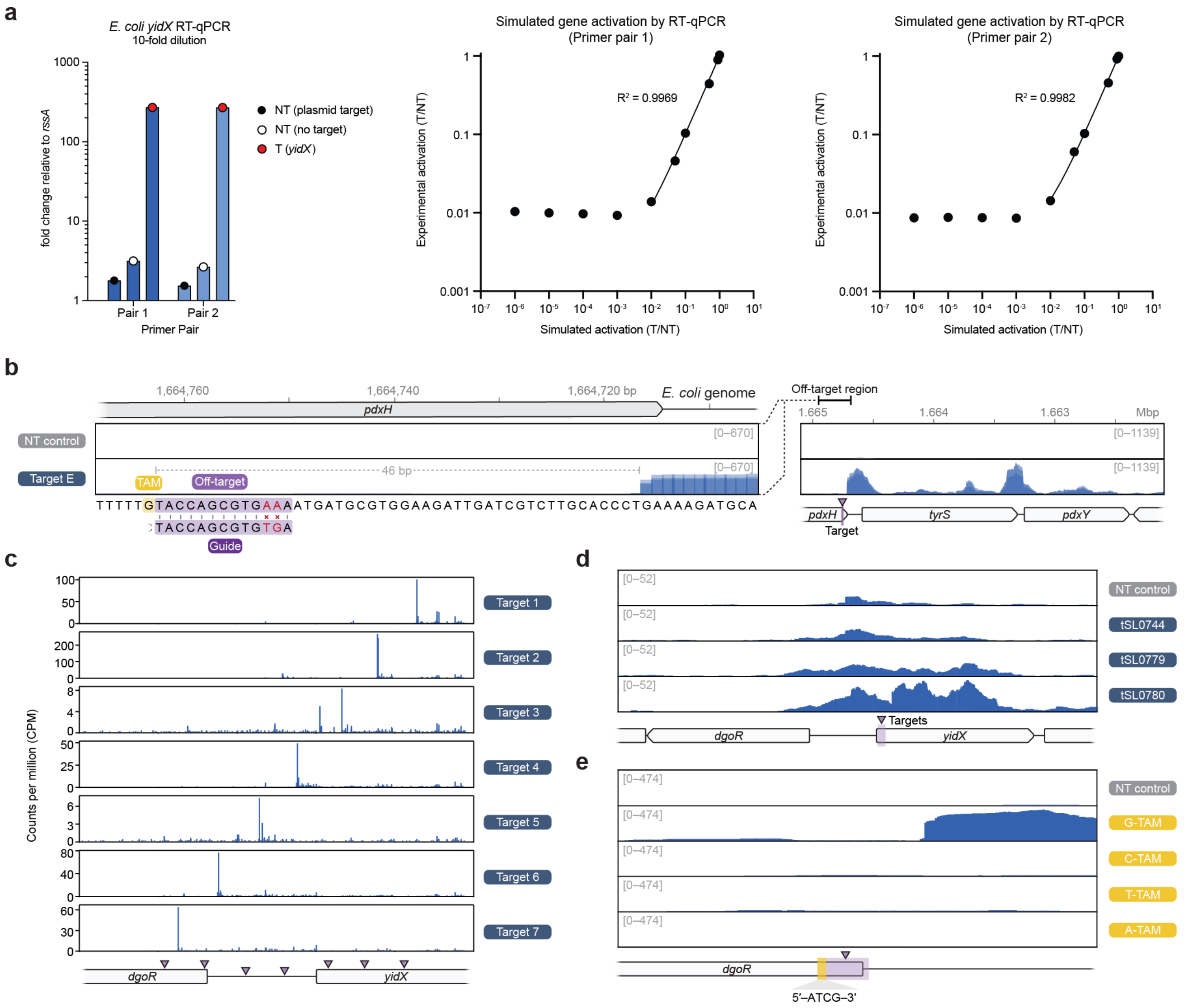


**Extended Data Figure 9 | Additional analyses relating to RNA-seq data. a,** RNA enrichment measured by reverse transcription (RT)-qPCR of target-E RNA-seq sample (**Fig. 6d**) for two distinct primer pairs (left). The targeting condition (T) reveals around 140-fold enrichment of transcripts versus non-targeting (NT) condition. Standard curves for simulated gene activation to determine the limit of detection for primer pair 1 (center) and primer pair 2 (right) suggest a detection limit of around 5%. **b,** RNA-seq coverage plots for the target-E off-target site near *tyrS*, shown in **Fig. 5f**. The ~46 bp distance between TAM and TSS upstream of *tyrS* is indicated. Zoom-out view (right) shows *pdxY* is upregulated due to it being encoded directly downstream of *tyrS*. The TAM and extensive 11-bp complementarity between the off-target DNA site and guide sequence are shown below the coverage track. Coverage is shown for the reverse strand. **c,** TSS plots derived from RNA-seq data, as shown in **Fig. 5c**, for all the individual gRNAs in **Fig. 5g**. **d,** RNA-seq tracks for a NT control and three distinct gRNAs designed for target-3 showing weak or no transcription initiation, likely due to binding site occlusion by other DNA binding factors involved in *yidX* regulation. Coverage is shown for the forward strand. NT, non-targeting. **e,** RNA-seq coverage plots for a gRNA designed for Target 6 (as shown in **Fig. 5g**) and three additional gRNAs that incrementally shift the TAM by 1 bp, changing it to A, T, and C. Targets flanked by a non-G-TAM fail to initiate transcription. Coverage is shown for the forward strand.

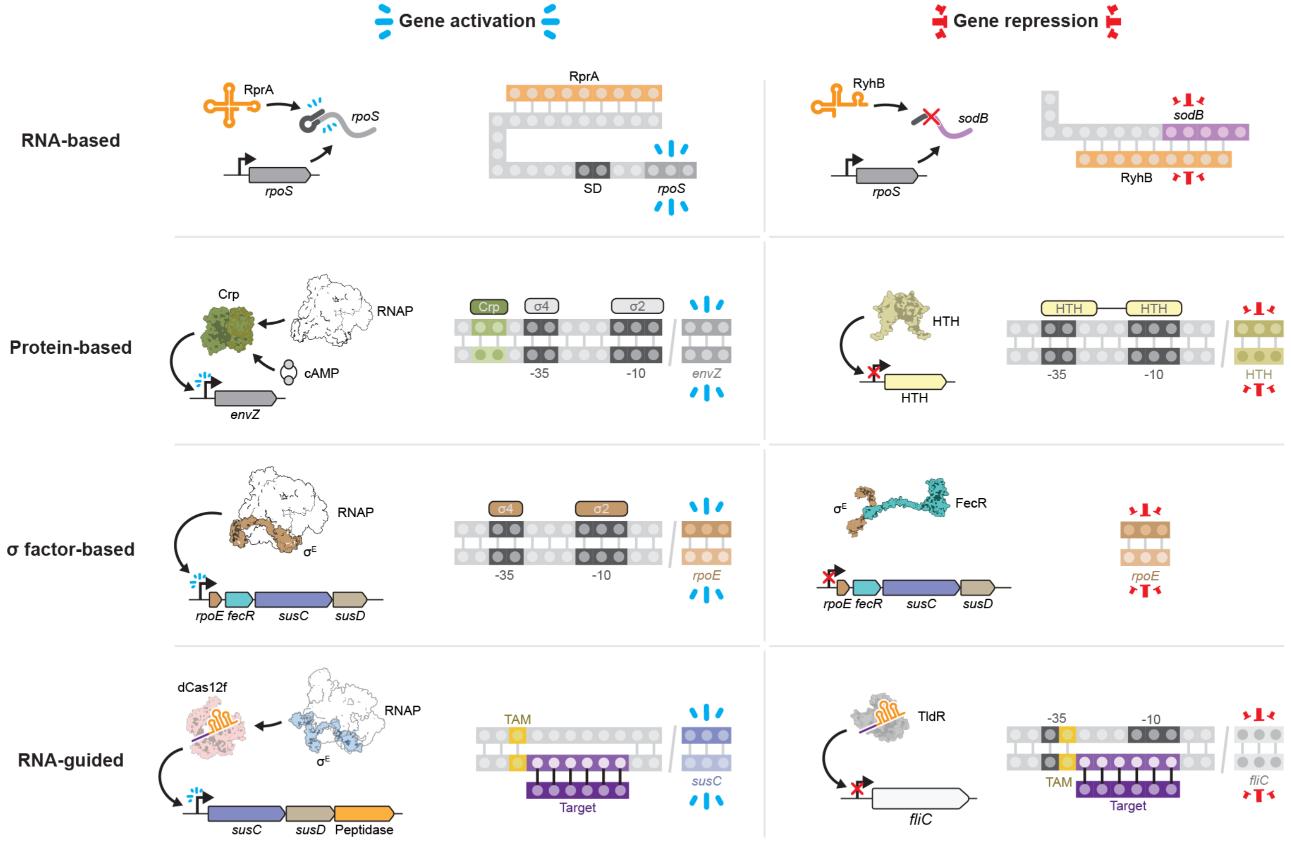


**Extended Data Figure 10 | Mechanisms of bacterial gene activation and repression, including newly discovered, RNA-guided pathways driven by dCas12f and TldR.** RNA-based mechanisms (top row) can activate gene expression, such as the RprA small RNA (sRNA) activating *rpoS* by relieving secondary structure inhibition, or repress gene expression, such as the RyhB sRNA destabilizing *sodB* through base pairing and RNase recruitment. Protein-based mechanisms (second row from top) can activate gene expression, such as Crp (cAMP receptor protein) enhancing RNAP recruitment at the *envZ* promoter, or repress gene expression, such as HTH transcription factors binding DNA to block RNAP-promoter recognition. σ factor-based mechanisms (second row from bottom) can activate gene expression, such as extracytoplasmic function (ECF) $\sigma^E$ factors recruiting RNAP to specific promoters to drive transcription, or repress gene expression, such as when their activities are inhibited by FecR anti-σ factors. Finally, we report novel RNA-guided pathways of gene regulation (bottom row). In previous work, we uncovered TnpB-like nuclease-dead repressors (TldR) that exploit gRNAs to bind complementary DNA target sites, thereby preventing promoter recognition by RNAP (bottom right). In this study, we uncover nuclease-dead Cas12f proteins that exploit gRNAs to bind complementary DNA target sites and directly recruit $\sigma^E$ factors and RNAP, thereby driving promoter-independent transcription of diverse genetic operons such as *susCD* polysaccharide utilization loci (bottom left). Collectively, our work highlights a new axis of gene regulation control via exapted, RNA-guided transcription factors akin to CRISPRi and CRISPRa.

### SUPPLEMENTARY FIGURES

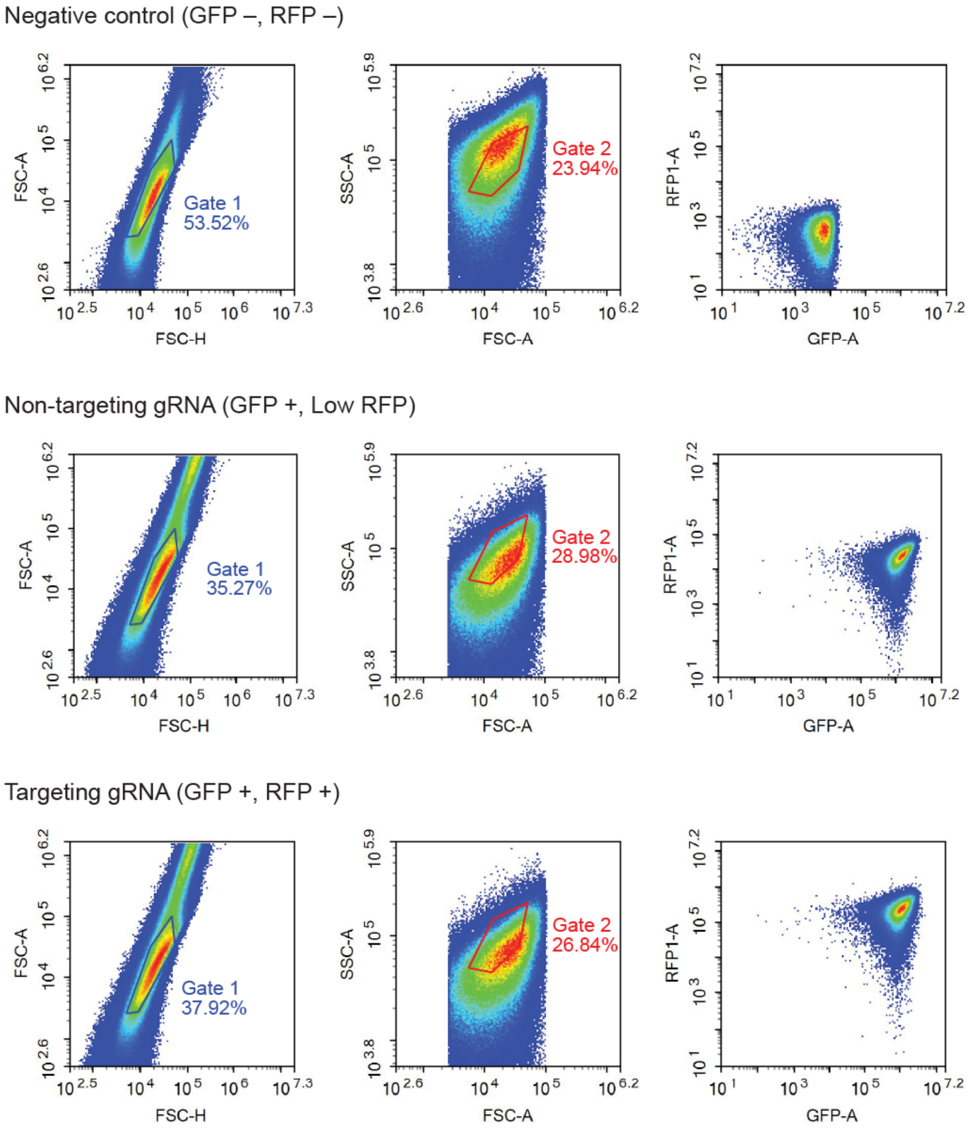


**Supplementary Figure 1 | Representative gating schemes for flow cytometry analysis of RFP fluorescence reporter.** Three representative samples are shown from top to bottom representing a strain with no fluorescence (Negative control, sSL0075), with a non-targeting gRNA (sSL4877; tSL0679), and an RFP-targeting gRNA (sSL4877; tSL0732) (**Supplementary Table 5**). Gates 1 and 2 separate single cells from doublets and debris. The fluorescence intensity is then measured for the final population in all samples and recorded for downstream analysis.
