## Supplementary Information Guide for "Exapted CRISPR-Cas12f homologs drive RNA-guided transcription"

### Supplementary Figures

Supplementary Figure 1 | Representative gating schemes for flow cytometry analysis of RFP fluorescence reporter.

### Supplementary Tables

Supplementary Table 1 | Cas12f homologs in Fig. 1b phylogenetic tree.

Supplementary Table 2 | dCas12f, HTH, and $\sigma^E$ homologs in Fig. 1e phylogenetic trees.

Supplementary Table 3 | $\sigma^E$ homologs in Fig. 1f phylogenetic tree.

Supplementary Table 4 | List of dCas12f-$\sigma^E$ systems, proteins, and gRNAs tested in this study.

Supplementary Table 5 | Strains used in this study.

Supplementary Table 6 | Description and sequence of plasmids used in this study.

Supplementary Table 7 | Oligonucleotides used in this study.

Supplementary Table 8 | Sequencing files and information for this study.
