## Supplementary Figure 1 for "Exapted CRISPR-Cas12f homologs drive RNA-guided transcription"

**Supplementary Figure 1 | Representative gating schemes for flow cytometry analysis of RFP fluorescence reporter.**

Three representative samples are shown from top to bottom representing a strain with no fluorescence (Negative control, sSL0075), with a non-targeting gRNA (sSL4877; tSL0679), and an RFP-targeting gRNA (sSL4877; tSL0732) (**Supplementary Table 5**). Gates 1 and 2 separate single cells from doublets and debris. The fluorescence intensity is then measured for the final population in all samples and recorded for downstream analysis.

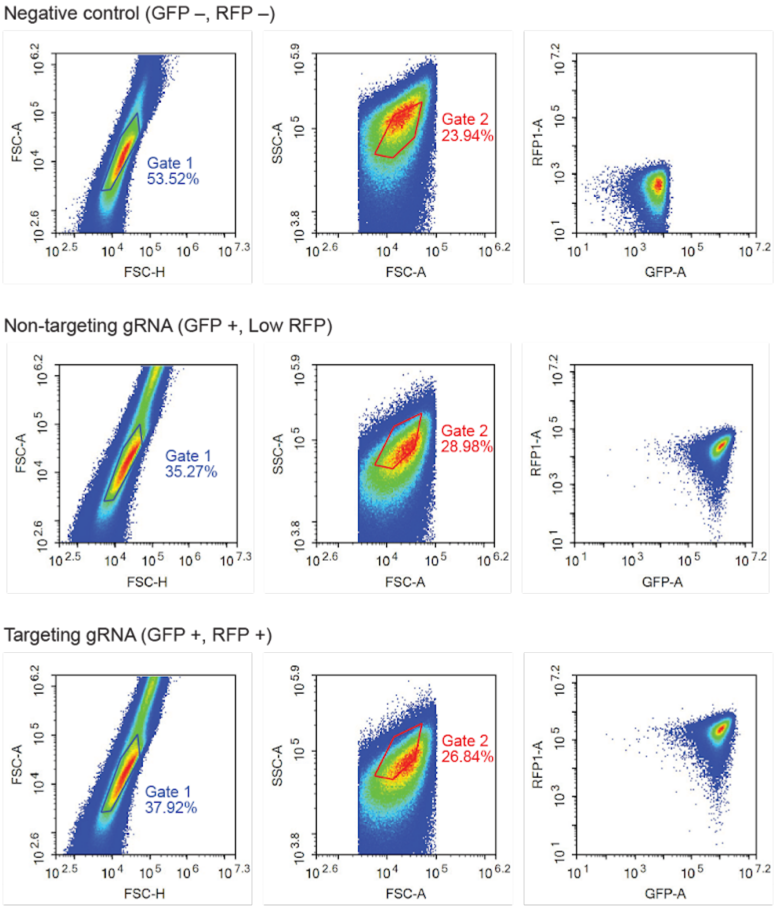
